# BAG3 regulation of Rab35 mediates the Endosomal Sorting Complexes Required for Transport /endolysosome pathway and tau clearance

**DOI:** 10.1101/2021.01.25.428055

**Authors:** Heng Lin, Maoping Tang, Changyi Ji, Peter Girardi, Gregor Cvetojevic, Yunpo Chen, Shon A. Koren, Gail V. W. Johnson

## Abstract

**Background:** Declining proteostasis with aging contributes to increased susceptibility to neurodegenerative diseases, including Alzheimer’s disease (AD). Emerging studies implicate impairment of the endosome-lysosome pathway as a significant factor in the pathogenesis of these diseases. Our lab was the first to demonstrate that BAG3 regulates phosphorylated tau clearance. However, we did not fully define how BAG3 regulates endogenous tau proteostasis, especially in the early stages of disease progression.

**Methods:** Mass spectrometric analyses were performed to identify neuronal BAG3 interactors. Multiple biochemical assays were used to investigate the BAG3-HSP70-TBC1D10B-Rab35-Hrs regulatory networks. Live-cell imaging was used to study the dynamic of endosomal pathway. Immunohistochemistry and immunoblotting were performed in human AD brains and BAG3 overexpressed P301S tau transgenic mice.

**Results:** The primary group of neuronal BAG3 interactors identified are involved in the endocytic pathway. Among them were key regulators of small GTPases, such as the Rab35 GTPase activating protein, TBC1D10B. We demonstrated that a BAG3-HSP70-TBC1D10B complex attenuates the ability of TBC1D10B to inactivate Rab35. Thus, BAG3 interacts with TBC1D10B to support the activation of Rab35 and recruitment of Hrs, initiating ESCRT-mediated endosomal tau clearance. Further, TBC1D10B shows significantly less co-localization with BAG3 in AD brains than in age-matched controls. Overexpression of BAG3 in P301S tau transgenic mice increased the co-localization of phospho-tau with the ESCRT III protein CHMP2B and reduced the levels of the mutant human tau.

**Conclusion:** We identified a novel BAG3-TBC1D10B-Rab35 regulatory axis that modulates ESCRT-dependent protein degradation machinery and tau clearance. Dysregulation of BAG3 could contribute to the pathogenesis of AD.

## Introduction

Maintaining proteostasis is vital for neuronal function and healthy aging. A critical aspect of proteostasis is an efficient system that recognizes and clears damaged or unnecessary proteins. Disruption of proteome homeostasis is likely a significant contributing factor to the pathogenesis of neurodegenerative diseases that are characterized by the accumulation of aggregation prone proteins, such as Alzheimer’s disease (AD)(1, 2). One of the defining hallmarks of AD is the presence of intraneuronal aggregates of phosphorylated tau (3). The accumulation of tau, particularly in an oligomeric state, advance AD pathogenesis (4). There is increasing evidence that compromised degradative mechanisms, and in particular lysosome dysfunction, contribute to the accumulation of toxic tau species and other disease-relevant proteins (5, 6).

Degradative pathways that direct cargos to the lysosome include autophagosome-lysosome and endosome-lysosome pathways, both of which are tightly regulated, vacuolar-based degradative systems. Although dysfunction of the autophagy pathway is likely one of the contributors to neurodegenerative proteopathies such as AD and Parkinson’s disease (7, 8), it is clear that impairment of the endosome-lysosome pathway is also a significant contributor to the pathogenesis of these diseases (9, 10). The endosome-lysosome pathway is an important protein clearance mechanism that directs the engulfment of protein cargo and trafficking to the lysosome for degradation. this process depends on the endosomal sorting complex required for transport (ESCRT) machinery, which mediates multivesicular body (MVB) formation and fusion with the lysosome (11). Deficits in ESCRT and the endosome-lysosome pathway, such as enlargement of the early endosome compartment and increased endosome/lysosome pH, are some of the earliest observable pathological changes in AD and other neurodegenerative diseases (12, 13). Further, cell models study demonstrated that compromised ESCRT leads to endolysosomal escape of tau seeds and propagation of tau aggregation (14). The mechanisms linking endosome-lysosome pathway dysregulation to AD pathogenesis have not been fully-elucidated..

Bcl-2-associated anthogene 3 (BAG3) is a stress-induced, multi-domain protein that plays a critical role in maintaining proteostasis, and thus neuronal health (15–18). As a stress-induced protein, BAG3 protein levels increase during aging (19). Interestingly, neurons with higher BAG3 expression are more resistant to tau pathology in AD (20), and recent data suggest that BAG3 levels are lower in AD brain compared to aged-matched controls (17). BAG3 promotes the clearance of tau (21) and other disease-relevant aggregation prone proteins such as α-synuclein (22), mutant SOD1, and mutant huntingtin (16). These findings point to the importance of BAG3 in maintaining neuronal proteostasis, which has historically been attributed to its role in autophagy, and this supposition is largely due to its ability to facilitate autophagy (15), although the mechanisms involved have not been fully defined and it is becoming apparent that this is likely not the only degradative pathway regulated by BAG3.

In the present study, we investigated the endogenous neuronal BAG3 interactome through unbiased immunoprecipitation-coupled mass spectrometry (IP-MS). Surprisingly, the endosome-lysosome pathway was the most over-represented. Of particular interest were regulators of small GTPases, such as the primary GTPase-activating protein (GAP) for Rab35, TBC1D10B ((Tre-2/USP6, BUB2, Cdc16) Domain Family 10B) (23). Rab35, which localizes to endosomes and the plasma membrane, participates in vacuole and protein trafficking, synapse vesicle turnover, and protein clearance (24–26). Activated Rab35 has been shown to recruit Hrs/ESCRT 0 and traffic tau as a cargo to the endosome (27), suggesting that Rab35 can regulate clearance of tau through the ESCRT-mediated endosome-lysosome pathway. However, the regulatory mechanisms underlying Rab35 in mediating tau clearance are not well understood.

In the present study, we demonstrate that the association of TBC1D10B with BAG3 disinhibits Rab35 activation, which in turn keeps Rab35 in a GTP bound, activated state resulting in enhanced ESCRT-0 recruitment and Hrs mobility. BAG3 also promotes tau recognition by ESCRT machinery, resulting in greater MVB trafficking of tau, lower hyperphosphorylated tau levels, and rescued neurite density in P301S tau (PS19 line) mice. Further, TBC1D10B shows significantly less co-localization with BAG3 in AD brains than in age-matched controls Our findings define a new role of BAG3 in regulating vacuolar-dependent protein degradation machinery through the TBC1D10B-Rab35-Hrs axis and provide a new therapeutic avenue for modulating tau clearance.

## Methods and Materials

### Reagents

Detail information of plasmids and antibodies can be found in Supplemental Methods and Materials in supplement 1.

### Animals

All animal procedures were approved by the University Committee on Animal Research of the University of Rochester. Detailed information of mouse and rat strains together with the sample preparation procedures are listed in Supplemental Methods and Materials in Supplemental Methods and Materials in supplement 1.

### Other experiment procedures

Experiment procedures including cell culture, generation of lentivirus, live cell imaging, stereotaxic surgeries, mass spectrometry analysis, Rab35 activity assay, immunoprecipitation, immunoblotting, immunohistochemical and Immunofluorescence staining can be found in supplement 1.

### Statistical analysis

All image measurements were obtained from the raw data. GraphPad Prism was used to plot graphs and perform statistical analysis. The statistical tests used are denoted in each Figure legend, and statistical significance was defined as *p < 0.05 and **p < 0.01. All immunoprecipitation data are repeated at least twice. For the live-cell imaging analysis, the difference of Hrs mobility curves (Figure 6d) was transformed into an accumulation curve and then analyzed using the Kolmogorov–Smirnov test.

## Results

Our previous findings, together with those from other groups, established BAG3’s role in autophagy (21, 28). However, as a co-chaperone protein with multiple domains, BAG3 likely has regulatory functions beyond autophagy through its interaction with other proteins. To better understand the role of BAG3 in mediating these processes, we coupled immunoprecipitation with mass-spectrometry (IP-MS) analysis of endogenous BAG3 from mature primary neuron cultures transduced with either scrambled or BAG3 shRNA. We identified 127 high-confidence BAG3-interacting proteins found enriched in the immunoprecipitates from neurons transduced with scrambled virus compared to shBAG3-mediated knockdown neurons (Supplement Table 1). We validated several of these putative BAG3 interactors, including MAP6 and clathrin heavy chain, by immunoprecipitation (Supplemental Figure S1A&B). KEGG analysis of these BAG3 interactors revealed endocytosis as the highest over-represented pathway (Figure 1A). Protein interaction mapping using STRING identified clusters of endocytosis-related and chaperone/co-chaperone proteins (Figure 1B and Supplemental Figure S2). In addition to chaperone and cytoskeletal proteins, other proteins such as GTPase-activating proteins (GAPs) play crucial roles in endocytosis (29, 30). TBC1D10B (also known as EPI64B or FP2461), is the major GAP for Rab35 (23) and was identified as a BAG3 interactor. Rab35 regulates endocytosis, recycling of synaptic vesicles, neurite elongation (24, 31), and the targeting of tau to the endolysosome compartment (27). Given the pivotal role of Rab35 in mediating vacuolar processes, neuronal health, and tau clearance, we investigated the potential role of BAG3 in mediating endosome/lysosome pathways via interacting with TBC1D10B.

**Figure 1.**
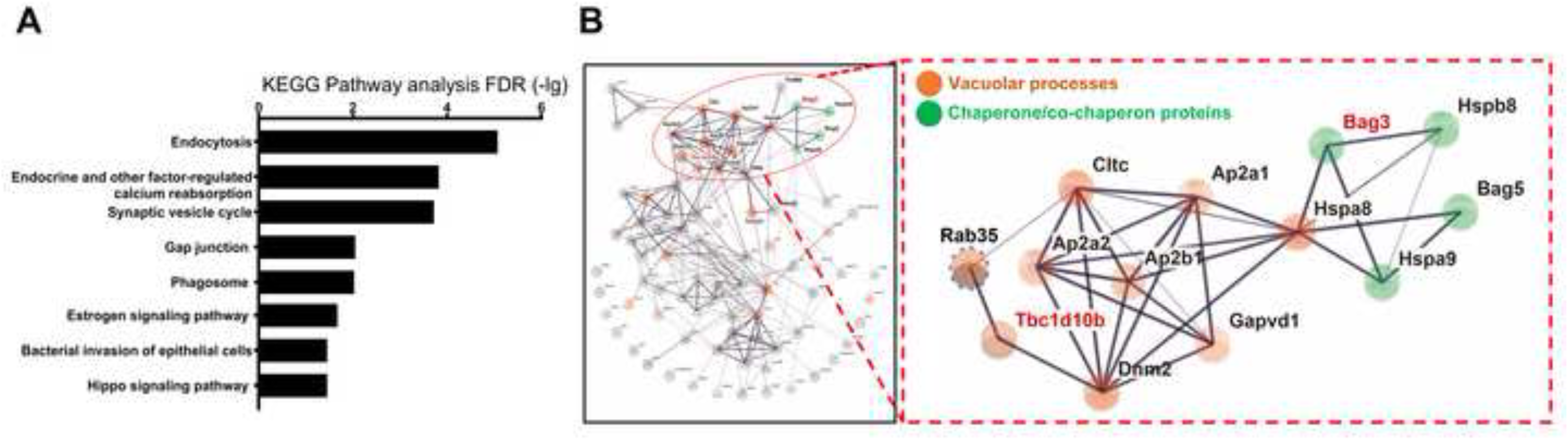
BAG3 interacts with proteins of the endocytosis pathway. Rat cortical neurons were transduced with lentivirus expressing scrambled (Scr) or BAG3 shRNA. Corresponding lysates were immunoprecipitated for BAG3 and associating proteins were run through LC-MS/MS. (A) KEGG enrichment analysis of BAG3 associated proteins with PSM ratios greater than 3 (Scr/shBAG3), FDR-adj. p-value < 0.1. (B) BAG3-associated protein interaction map using STRING. Line thickness indicates the literature interaction. Nodes directly linked to vacuolar processes are labeled in red, and nodes that are related to chaperone/co-chaperone proteins are labeled in green.

### BAG3 interacts with TBC1D10B in hippocampal and cortical neurons

To verify the interaction of TBC1D10B with BAG3, we immunoprecipitated TBC1D10B from rat cortical neuron lysates and immunoblotted for BAG3. Endogenous BAG3 readily co-precipitated with TBC1D10B (Figure 2A). We next investigated the spatial localization of BAG3 and TBC1D10B in mouse brain. Immunohistochemistry (IHC) of 8-month-old wildtype mouse brain showed TBC1D10B puncta co-localized with BAG3 throughout neurons in the CA1 region of the hippocampus (Figure 2B). We further examined the co-localization of BAG3 with TBC1D10B in cultured rat cortical neurons. Like our *in vivo* observations, TBC1D10B co-localized with BAG3 in the neurites and the soma in punctate-like structures (Figure 2C-G). Co-localization analysis (32) showed overlap between TBC1D10B puncta and BAG3 in the soma (12.4 ± 2.8 %, mean ± SEM) and neurites (8.0 ± 2.0 %). Together, these data show BAG3 and TBC1D10B interact and co-localize in neurons. Extending this to postmortem human brains, the co-localization of BAG3 with TBC1D10B was significantly reduced in human AD brain samples compared with age matched controls (Figure 2H-K).

**Figure 2.**
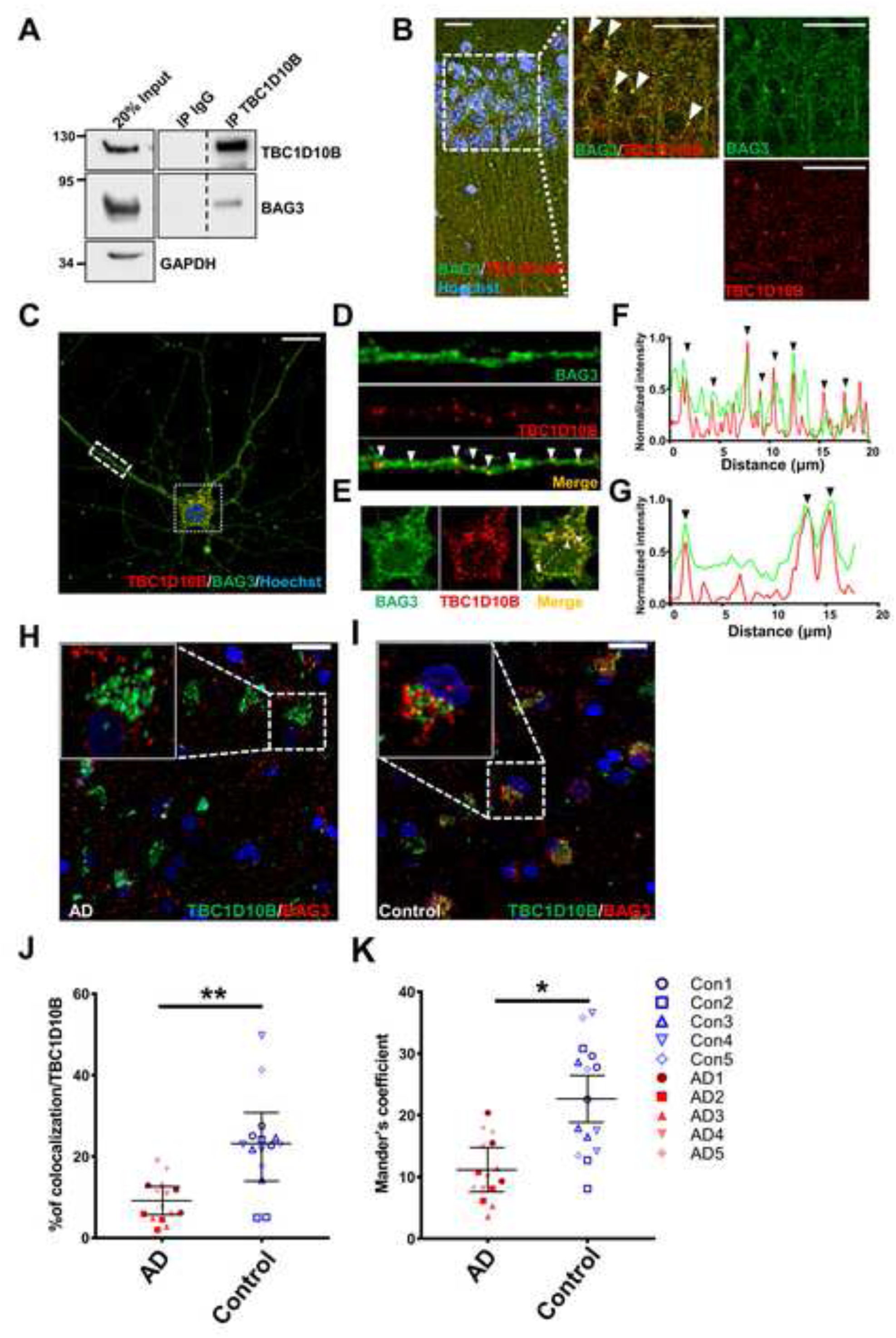
BAG3 associates with TBC1D10B in neurons. (A) Rat cortical neuron lysates were immunoprecipitated with an anti-TBC1D10B antibody and immunoblotted for BAG3. An indicated fraction of cell lysate was used as input control with GAPDH as loading control. Immunoprecipitates were probed for TBC1D10B, BAG3, and Rab35. Vertical dashed lines indicate that intervening lanes were removed. However, all images were from the same blot and exposure. (B) Representative immunostaining of BAG3 (green) and TBC1D10B (red) co-localization in the CA1, counterstained with Hoechst 33342. Scale bars, 20μm. (C-G) Neurons were immunostained for BAG3 (red) and TBC1D10B (green). Overlap of BAG3 with TBC1D10B puncta was observed in the neurites (D) and soma (F) with corresponding line scans (E&G). Arrowheads indicate areas of overlap. Scale bar, 10 μm.

### The interaction of BAG3 with TBC1D10B is facilitated by HSP70

Given the plethora of domain-specific BAG3 interactors (16), we next assessed which domain in BAG3 facilitate TBC1D10B interaction using a co-expression model in BAG3 knockout (KO) HEK cells. FLAG-tagged TBC1D10B was co-expressed with wildtype (WT) and mutant forms of BAG3 followed by co-immunoprecipitation analysis (Figure 3A). As predicted by the co-localization analysis, WT BAG3 readily associates with TBC1D10B and not in BAG3 KO cells. The WAWA mutant form of BAG3, which prevents BAG3 WW domain association with synaptopodin-2 (myopodin) (33) and synaptopodin (28), decreased the apparent co-immunoprecipitation of BAG3 with TBC1D10B. The GPG mutation in BAG3, which is a mutation of the IPV domain that prevents association with small heat shock proteins (34, 35), did not alter association with TBC1D10B. Strikingly, the L462P mutation in BAG3, which is in the C-terminal BAG domain which mediates binding with HSP70 (36, 37), abolished its interaction with TBC1D10B. L462P is a rare mutation in BAG3 that causes cardiomyocyte protein aggregation and dilated cardiomyopathy (37) and has been speculated to decrease BAG3-HSP70 association. To assess whether HSP70 is involved in modulating BAG3-TBC1D10B interaction, we first verified that the L462P mutation of BAG3 strongly decreased (>50%) the association with HSP70 using a co-expression model of WT or L462P BAG3 with V5-HSP70 in BAG3 KO cells (Figure 3B). Next, we examined whether modulating HSP70 levels or activity regulated BAG3-TBC1D10B interaction. HSP70 over-expression resulted in enhanced BAG3 co-immunoprecipitation with TBC1D10B compared to mock-transfected HEK cells (Figure 3C). Pharmacological inhibition of HSP70 ATPase activity and HSP70-BAG3 association using 10µm YM-01 (38) disrupted BAG3-TBC1D10B co-immunoprecipitation (Figure 3D). Pairing V5-HSP70 over-expression with YM-01 treatment prevented this loss of BAG3-TBC1D10B association, likely due to saturation of YM-01 with exogenous HSP70. Overall, these results suggest HSP70 facilitates the association of BAG3 with TBC1D10B.

**Figure 3.**
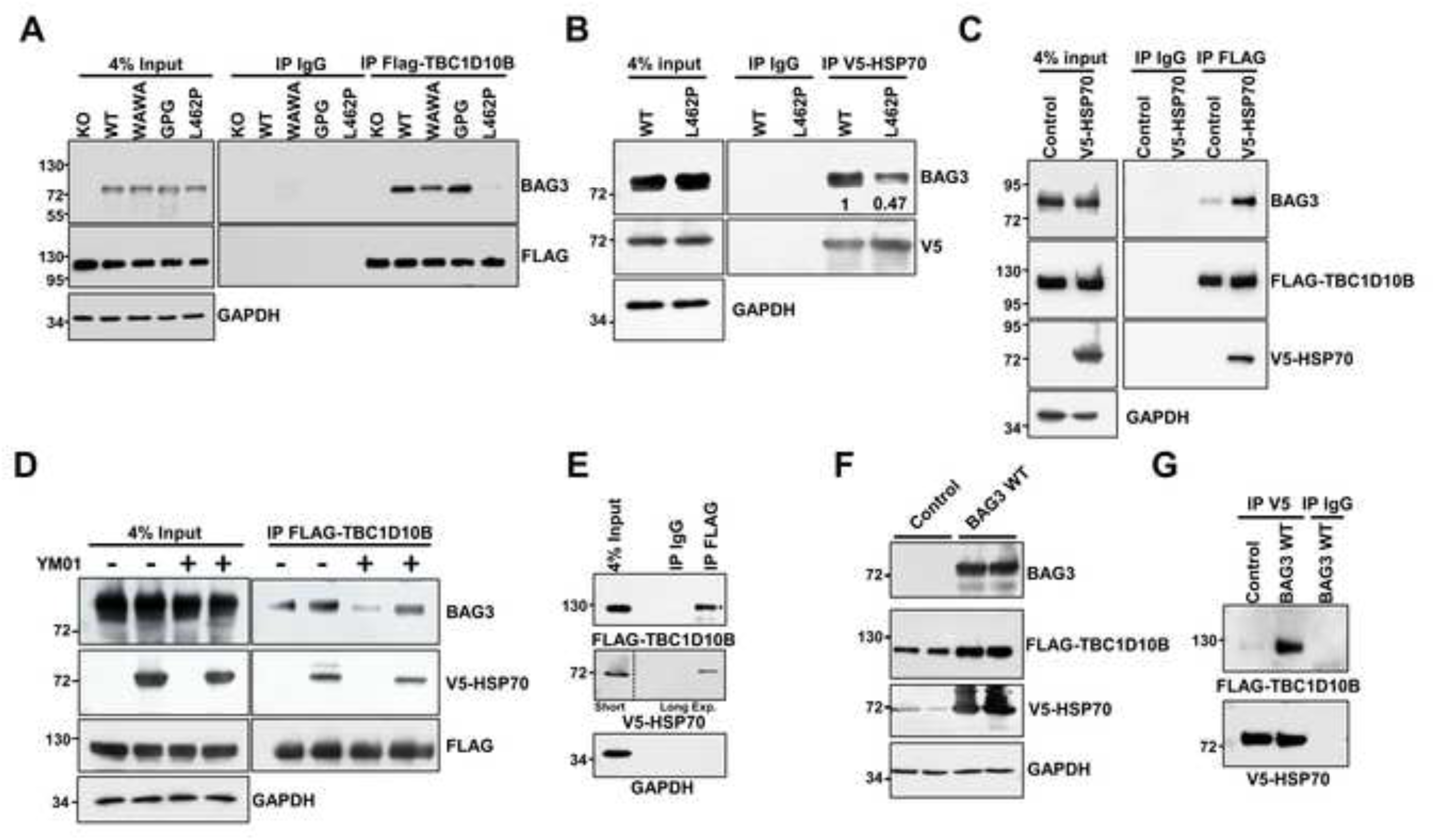
HSP70 facilitates the association of BAG3 with TBC1D10B at the BAG domain. (A) BAG3 null HEK293TN cells were transiently transfected with FLAG-TBC1D10B together with empty vector (Control), wild type BAG3 (WT BAG3), or indicated BAG3 mutants. Cell lysates were immunoprecipitated with an anti-FLAG antibody, followed by blotting for BAG3 and FLAG. Immunoprecipitates were probed for FLAG and BAG3. (B) BAG3 null HEK293TN cells were transiently transfected with V5-HSP70 together with wild type BAG3 (WT) or L462P BAG3 (L462P). Cell lysates were immunoprecipitated with a V5 tag antibody, followed by blotting for BAG3 and V5. (C) HEK293TN cells were transiently transfected with FLAG-TBC1D10B together with an empty vector (Control) or V5-HSP70. Cell lysates were immunoprecipitated with an anti-FLAG antibody, followed by blotting for endogenous BAG3, FLAG, and V5. (D) HEK293TN cells were transient co-expressed with FLAG-TBC1D10B together with an empty vector or V5-HSP70 and treated with 10 μM YM01 or DMSO (vehicle control). Corresponding lysates were collected and immunoprecipitated with anti-FLAG antibody, followed by blotting for BAG3, V5, and FLAG. (E) BAG3 null HEK293TN cells were transiently transfected with V5-HSP70 and FLAG-TBC1D10B. Corresponding lysates were immunoprecipitated with anti-FLAG antibody, followed by blotting for V5 and FLAG. Results of different exposure times for the same blot were separate with a vertical dotted line. Indicated fraction of cell lysate was used as input control. (F) BAG3 null HEK293TN cells were transiently transfected with FLAG-TBC1D10B, V5-HSP70 together with an empty vector (Control) or wildtype BAG3 (BAG3 WT). Cell lysates were immunoprecipitated with anti-FLAG antibody, followed by blotting for BAG3, V5, and FLAG. (G) Cell lysate from F was immunoprecipitation with the 0.5µg IgG or V5 antibody which is saturated by V5-HSP70 and blotted for FLAG. GAPDH was used as a loading control for all input lanes.

These findings prompted us to examine if HSP70 associated directly with TBC1D10B, independent of BAG3. Co-expressing FLAG-TBC1D10B and V5-HSP70 in BAG3 KO HEK cells followed by immunoprecipitation with V5 antibody revealed HSP70 associated with TBC1D10B in the absence of BAG3 (Figure 3E). BAG3 interaction prevents the degradation of the small heat shock protein HspB8 (35, 39) and we identified that the same is true for HSP70, as increased expression of BAG3 greatly increases HSP70 levels (Figure 3F). To examine if BAG3 could also regulate the association between HSP70 with TBC1D10B, we co-transfected V5-HSP70, FLAG-TBC1D10B, and WT BAG3 or empty vector followed by a saturated immunoprecipitation with the V5 antibody. This resulted in equal amounts of V5-HSP70 in the control and BAG3 overexpression group pulled down with V5 (Figure 3G). These data demonstrated that BAG3 greatly enhances the association of HSP70 with TBC1D10B. Overall, our data indicate that HSP70 facilitates the association of BAG3 and TBC1D10B, and that HSP70, TBC1D10B and BAG3 interact to form a complex.

### BAG3 regulates the Rab35 activity through the association of TBC1D10B

TBC1D10B is a GAP that specifically associates with the GTP bound, active form of Rab35 and promotes GTP hydrolysis, leading to the inactivation of Rab35 (23). Since our data showed that BAG3 associates with TBC1D10B, we examined how BAG3 may regulate the function of TBC1D10B. Rat cortical neurons were transduced with lentivirus expressing scrambled or BAG3 shRNA, followed by immunoprecipitation of TBC1D10B and immunoblotting for Rab35. Our data showed neuronal BAG3 knockdown alters TBC1D10B-Rab35 association without modifying total levels of TBC1D10B (Figure 4A). This finding leads to two opposing hypotheses: 1) Because GAPs preferentially bind GTP bound Rabs (40, 41), BAG3 promotes the association of TBC1D10B with Rab35 and leads to the inactivation of Rab35 (conversion to GDP-Rab35) and thus decreases Rab35-TBC1D10B interaction; 2) Binding of BAG3 attenuates the ability of TBC1D10B to stimulate the GTPase activity of Rab35, resulting in prolonged association of TBC1D10B with Rab35 (42). To test these hypotheses, we used an established Rab35 activity assay based on GST-tagged RBD35, which specifically binds the active, GTP-bound form of Rab35 (43). Cell lysates from rat neurons transduced with lentivirus expressing scrambled or BAG3 shRNA were incubated with GST or GST-RBD35 on glutathione beads, and the precipitates blotted for Rab35. Our data show that knockdown of BAG3 did not alter total protein levels of Rab35, but instead reduced GTP-bound Rab35 (Figure 4B). This finding supports our second hypothesis that BAG3 keeps Rab35 in a GTP-bound state by associating with TBC1D10B, thereby preventing Rab35 inactivation.

**Figure 4.**
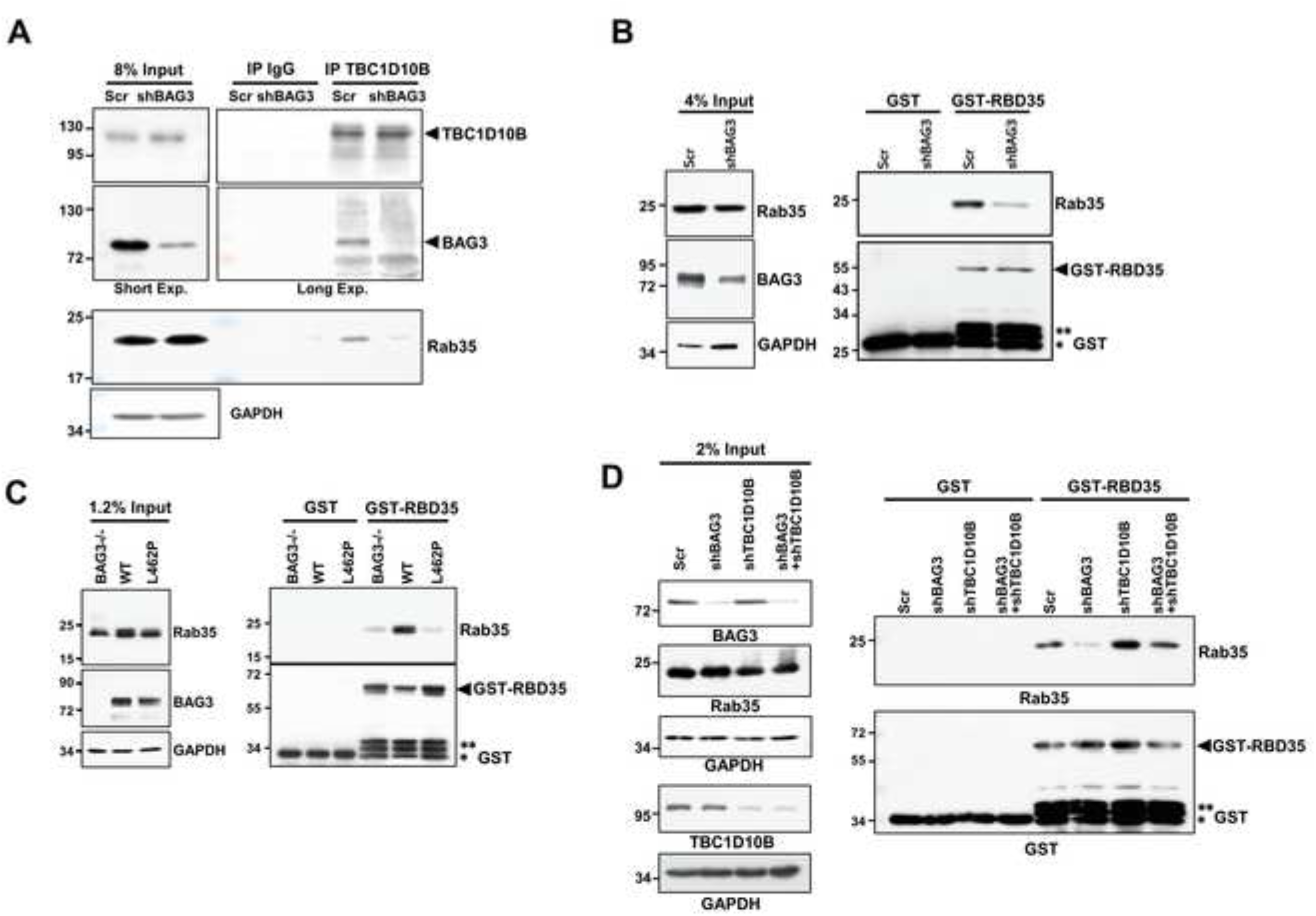
BAG3 associates with TBC1D10B to regulate the Rab35 activity. (A) Rat cortical neurons were transduced with lentivirus expressing shBAG3 or a scrambled (Scr) version. Cell lysates were immunoprecipitated with an anti-TBC1D10B antibody and immunoblotted for BAG3, TBC1D10B, and Rab35. An 8% fraction of cell lysate was used as input control. (B) Rat cortical neurons were transduced with lentivirus expressing scrambled (Scr) or shBAG3 shRNA. Rab35 activity was examined by incubation of cell lysates with purified GST or GST-RBD35, followed by precipitation with glutathione beads (43). The precipitated samples were blotted for Rab35 and GST. 4% of the cell lysate was used as input control. (C) BAG3 null HEK293TN cells were transiently transfected with Myc-Rab35 together with empty vector (Control), wild type BAG3 (WT BAG3) or L462P BAG3. Rab35 activity was examined by pull-down with GST-RBD35, as described in B. The precipitated samples were blot for Rab35 and GST. 1.2% of the cell lysate was used as input control. (D) Rat cortical neurons were transduced with lentivirus expressing scrambled (Scr), shBAG3, shTBC1D10B, or both shBAG3 and shTBC1D10B shRNAs. Rab35 activity was examined by pull-down with GST-RBD35. Arrow indicates the GST-RBD35 fusion protein; **, GST-RBD35 degradation products, *, GST protein. GAPDH was used as a loading control for all input lanes.

To examine if disrupting the association of BAG3 with TBC1D10B could also affect Rab35 activity, we co-transfected Myc-Rab35 with WT BAG3, L462P BAG3, or empty vector in BAG3 KO HEK cells. Repeating the Rab35 activity assay with these conditions showed that loss of BAG3 substantially decreased the level of GTP-bound Rab35 without affecting its expression level, consistent with our findings in rat cortical neurons (Figure 4C). Interestingly, the L462P mutation of BAG3, which greatly disrupted BAG3 association with TBC1D10B, also decreased the level of GTP-bound Rab35 (Figure 4C).

Our findings suggest that BAG3 associates with TBC1D10B in order to maintain Rab35 in an active state. In contrast, depletion of BAG3 or freeing TBC1D10B from BAG3 will increase the inactivation of Rab35. This finding further leads to another hypothesis that a primary mechanism by which BAG3 regulates the Rab35 activity is through mediating the function of TBC1D10B. To test whether BAG3 regulates Rab35 activity through TBC1D10B, we examined Rab35 activity in conditions with BAG3, TBC1D10B, or both were knocked down using shRNA. Resulting data demonstrated that, compared to scrambled shRNA conditions, GTP-bound Rab35 levels are increased when TBC1D10B is knocked down, but decreased when BAG3 is knocked down. The decreased GTP-bound Rab35 levels observed with BAG3 knockdown was alleviated when TBC1D10B was knocked down as well (Figure 4D). Our findings suggest a model where BAG3 inhibits the function of TBC1D10B, maintaining active Rab35 levels. Conditions with depleted BAG3, such as in AD(17), could therefore lead to inactivated Rab35 and impaired endocytosis.

### BAG3 interacts with TBC1D10B to regulate tau sorting into the endocytic pathway through the ESCRT system

Rab35 is a GTPase that plays an essential role in the endocytic pathway and facilitates tau clearance in neurons (27). As shown in primary rat cortical neurons, BAG3 promotes tau degradation (21) and interacts with TBC1D10B (Figure 2A). BAG3 TBC1D10B association ultimately regulates the activity of Rab35 (Figure 4B), which leads to TBC1D10B potentially playing a role in phosphorylated tau clearance as well. Indeed, our data shows that depletion of TBC1D10B in neurons significantly decreased the levels of p-Thr231, p-Ser262, and p-Ser396/Ser404 tau in mature neurons (Figure 5A&B). We next examined whether combining knockdown of TBC1D10B and BAG3 rescued phosphorylated tau levels (Figure 5C). First, we supported our previous results showing BAG3 knockdown increased the levels of p-Thr231, p-Ser262, and p-Ser396/Ser404 tau in mature neurons (21). Consistent with our proposed model, simultaneous depletion of TBC1D10B and BAG3 negated this effect on p-tau levels compared with BAG3 knockdown alone (Figure 5D&E). These data suggested that TBC1D10B is downstream of BAG3 in regulating tau clearance. Since Rab35 clears tau through the ESCRT system (27), we hypothesized that BAG3 acts in concert with TBC1D10B to regulate phosphorylated tau clearance by similar means. We assessed whether BAG3 knockdown altered the co-localization between CHMP2B, an ESCRT III component (44), and p-Ser396/404 tau in neuronal processes. In BAG3 knockdown neurons, CHMP2B colocalized with p-Ser396/404 tau significantly less compared with the scrambled group. Depletion of both BAG3 and TBC1D10B significantly increased the co-localization of p-Ser396/404 tau with CHMP2B (Figure 6), indicating that BAG3 acts with TBC1D10B to regulate phosphorylated tau clearance through the ESCRT system.

**Figure 5.**
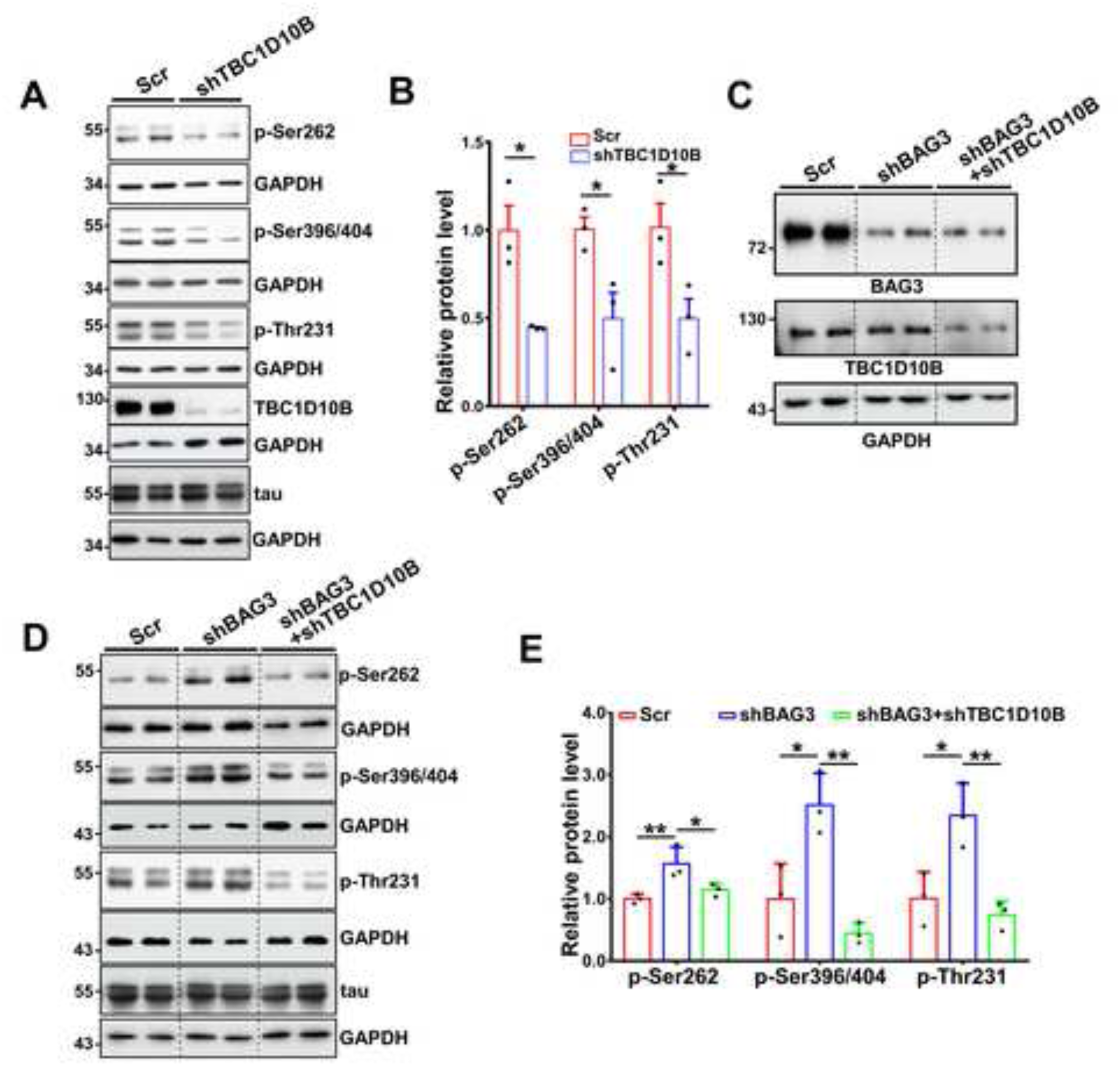
BAG3 cooperates with TBC1D10B to regulate tau levels in neurons. (A) Rat cortical neurons were transduced with lentivirus expressing scrambled (Scr) or shTBC1D10B shRNA. Cell lysates were immunoblotted for TBC1D10B, p-Ser262, p-Ser396/404, and p-Thr231 tau. GAPDH is used as a loading control. (B) Relative level of p-tau to total tau. Data are shown as mean ± SEM with unpaired Student’s t-test; *, P<0.05, **, P<0.01. (C-D) Rat cortical neurons were transduced with lentivirus expressing scrambled (Scr) or shBAG3 or both shBAG3 and shTBC1D10B shRNAs. Cell lysates were either immunoblotted for BAG3 and TBC1D10B for validating the knockdown (C) or p-Ser262, p-Ser396/404, and p-Thr231 tau (D). (E) Relative level of p-tau to total tau. Data are shown as mean ± SEM with one-way ANVOA and Tukey’s multiple comparisons test. *, P<0.05, **, P<0.01. For all experiment n=3 for each group. Vertical dotted lines indicate that intervening lanes were removed, however, all images were from the same blot and exposure.

**Figure 6.**
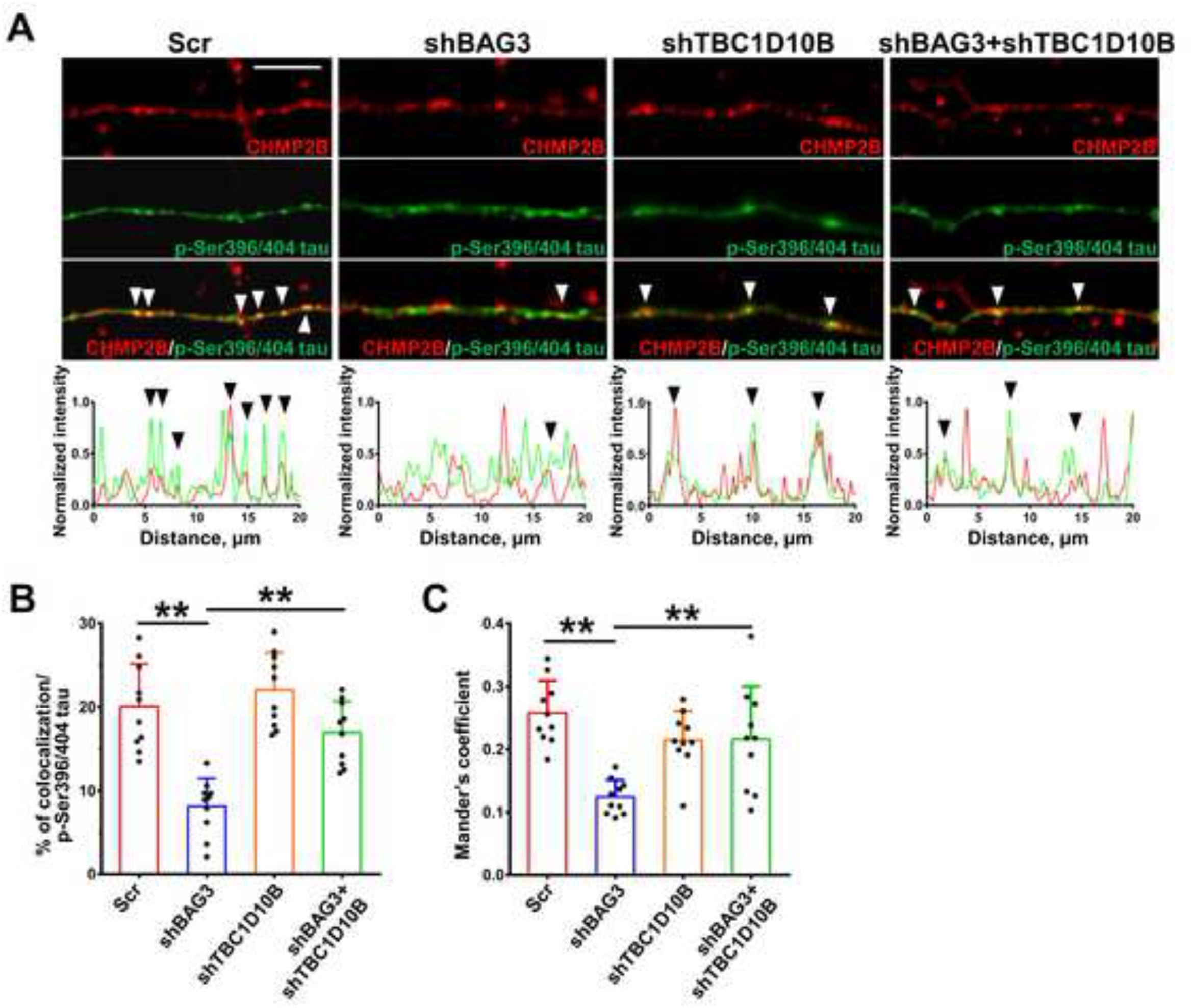
BAG3 interacts with TBC1D10B to regulate the tau sorting into the endocytic pathway through the ESCRT system. (A) Rat cortical neurons were transduced with lentivirus expressing scrambled (Scr), shBAG3, shTBC1D10B or both shBAG3 and shTBC1D10B shRNAs. Neurons were co-immunostained for CHMP2B (red) and p-Ser396/404 tau (green). The corresponding line scans are shown below for neurites. Arrowheads indicate areas of overlap. Scale bar, 5 μm. (B) Quantification of the co-localization between CHMP2B and p-Ser396/404 tau based on volume. (C) Quantification of co-occurring of CHMP2B in p-Ser396/404 tau using Mander’s coefficient. Data were presented as mean ± SD, N=10 neurites from 6+ cells each with one-way ANVOA and Tukey’s multiple comparisons test. *, P<0.05, **, P<0.01.

### BAG3 regulates the recruitment of Hrs to Rab35

Rab35 plays an essential role in the ESCRT system. When activated, Rab35 recruits Hrs, an ESCRT 0 component, to the surface of the endosome to facilitate the recruitment of other ESCRT machinery to form intraluminal vesicles and in directing cargo, such as phospho-tau species (27), to the endosome for engulfment (31). Since our data suggest that BAG3 regulates the activity of Rab35 through interacting with TBC1D10B, we hypothesize that BAG3, by promoting active Rab35, should facilitate the recruitment of Hrs to Rab35. Therefore, we investigated the co-localization of Rab35 with Hrs in primary rat cortical neurons. We found Rab35 co-localizes with Hrs both in the soma and neuronal processes, and that this co-localization was significantly decreased in BAG3 knockdown neurons (Figure 7A-F). The recruitment of Hrs to Rab35 to initiate the ESCRT pathway is a dynamic process, in that Hrs must be both recruited and released for intraluminal vesicle formation of MVBs (45). We next examined Rab35 and Hrs dynamics using live cell imaging of co-transfected eGFP-Rab35 and Hrs-RFP with and without BAG3. In BAG3 KO HEK cells, the majority of Hrs puncta were stationary and only 4.5% of the Hrs puncta moved more than 1μm over 10 minutes (Figure 7G&H). In contrast, rescuing BAG3 expression with WT-BAG3 rescued Hrs puncta mobility, with 23% of the Hrs puncta moving more than 1μm over 10 minutes (Figure 7G&H). Additionally, we observed the co-localization of Rab35 with Hrs was greatly increased in the presence of BAG3 compared with the BAG3 KO condition (Figure 7I). These findings suggest BAG3 enhances the mobility of Hrs and boosts the recruitment dynamics of Hrs to Rab35. Next, we examined the regulatory function of BAG3 and TBC1D10B on the association of Hrs and Rab35 by expressing Myc-Rab35 with and without overexpression of BAG3 and TBC1D10B. Co-immunoprecipitation analysis revealed that Rab35 and Hrs minimally associate in BAG3 KO cells, but returned to a strong association when BAG3 was re-introduced (Figure 7J). Further, overexpression of TBC1D10B disrupts the association of Rab35 with Hrs in the presence of BAG3 (Figure 7J). These findings suggest BAG3 strongly promotes the recruitment of Hrs to Rab35, and TBC1D10B acts downstream of BAG3 to inhibit this recruitment.

**Figure 7.**
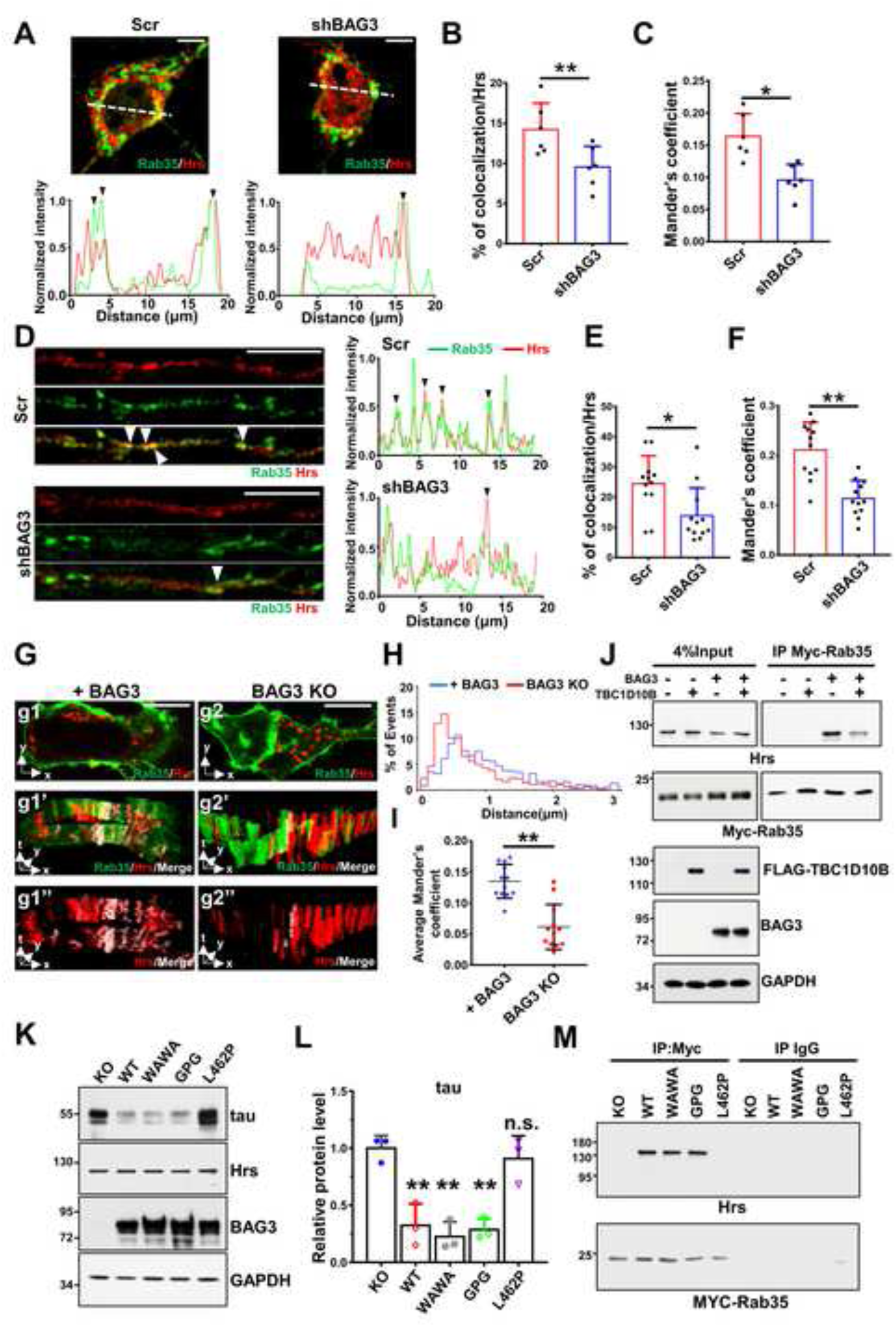
BAG3 and TBC1D10B regulate the mobility of Hrs and its interaction with Rab35. (A-F) Rat cortical neurons that were transduced with lentivirus expressing scrambled (Scr) or shBAG3 shRNAs were co-immunostained for Rab35(Green) and Hrs (Red). Overlap of Rab35 with Hrs puncta was observed in the soma (A) and neurites (D). Scale bars, 10 μm. The corresponding line scans are shown at the bottom for A or right for D. Arrowheads indicate areas of overlap. (B, E) Quantification of the co-localization between Rab35 and Hrs based on volume. (C, F) Quantification of co-occurring Rab35 in Hrs using Mander’s coefficient. N=12 neurites from 6 different neurons in each group. Data was presented as mean±SD with unpaired Student’s t-test., *P<0.05, **P<0.01. (G) BAG3 null HEK293TN cells were transfected with BAG3 WT (+BAG3) or empty vector (BAG3 KO), RFP-Hrs, and GFP-Rab35. Live-cell imaging was imaged at 0.1 Hz for 10 minutes. G1 and G2, are representative snapshots of HEKS with or without BAG3. G1’, G1’’ and G2’, G2’’ are 3D kymographs of G1 and G2. The white color indicates the co-localized region of Rab35 and Hrs over time. Scale bars, 20µm. (H) The graph shows the track distance of Hrs puncta in 10 minutes. The distance of the track was binned every 0.1 μm. The percentage of events is defined as the percentage of tracks of Hrs in a certain distance range in one cell. At least 10 cells from each group were analyzed. (p < 0.01, Kolmogorov-Smirnov test). (I) Comparison of co-occurring Rab35 in Hrs using Mander’s co-localization coefficient over time in BAG3 expressing and BAG3 KO cells. Mean±SD, N=12, with unpaired Student’s t-test. **, P<0.01. (J) BAG3 null HEK293TN cells were transfected with Myc-Rab35 and with/without BAG3 and TBC1D10B. Corresponding cell lysates were immunoprecipitated with anti-Myc antibody and immunoblotted for Hrs and Myc. Four percent of the lysates were used for input control. (K-M) BAG3 null HEK293TN cells stably expressing tau were transfected with Myc-Rab35 together with empty vector (Control), wildtype BAG3 (BAG3 WT), or BAG3 mutants. (K) Corresponding lysates were immunoblotted for total tau, Hrs and BAG3 and quantified in (L). N=3 for each group. Data are shown as mean ± SEM with one-way ANOVA and Tukey’s multiple comparisons test. **, P<0.01. n.s. not significant. (M) Cell lysates were immunoprecipitated with Myc antibody and immunoblotted for Hrs. For immunoblots shown in F, G, and I, GAPDH was used as a loading control.

As specific domains of BAG3 were involved in mediating its interaction with TBC1D10B, we hypothesized that different domain mutations of BAG3 would diminish the recruitment of Hrs to Rab35. Since Hrs recruitment is important to initiate the ESCRT pathway to sort tau into endosome-lysosome for degradation (27, 45, 46), we also hypothesize that mutations of BAG3 would decrease tau degradation in a domain-dependent manner. To test these hypotheses, we generated stable tau-expressing BAG3 KO HEKs cells by transducing BAG3 KO HEKs cells with 0N4R tau-T2A-RFP lentivirus, followed by fluorescence-activated cell sorting. We then co-transfected Hrs, Myc-Rab35, and WT BAG3 or BAG3 with different mutations into the BAG3 KO HEKs cells. As expected, WT BAG3, WAWA BAG3, and GPG BAG3 expression significantly reduced tau levels in BAG3 KO HEKs (Figure 7K&L). Expression of L462P BAG3 failed to decrease tau levels, further supporting the importance of BAG3-HSP70 interactions in tau degradation (Figure 7G&H). Next, we immunoprecipitated Myc-Rab35 and blotted for Hrs. We found that expressing WT BAG3, WAWA BAG3 or GPG BAG3 enhanced the association of Hrs with Rab35 compared with the BAG3 KO condition, while the L462P BAG3 expression group showed minimal association of Hrs with Rab35 (Figure 7M). Since the L462P mutation of BAG3 disrupted the association of BAG3 and TBC1D10B, our findings indicate that BAG3 interacts with TBC1D10B to regulate the Hrs recruitment to Rab35.

### Hippocampal overexpression of BAG3 alleviates the tau pathology development in P301S mice

Recent reports have suggested BAG3 maintains neuronal proteostasis and acts as a protector against tau aggregation (17, 19, 21). We tested whether BAG3 over-expression reduces tau pathology in sex-controlled P301S tau transgenic mice (PS19 line) (47–49). For these studies we only used male P301S as it has been reported that they develop tau pathology more consistently than females (48, 49). Two-month-old P301S mice were subjected to bilateral intrahippocampal AAV injections of T2A-hBAG3 (with an N-terminal FLAG-Myc tag) or eGFP control and collected at 6 months of age. Resulting IHC data showed endogenous BAG3 is expressed both in the soma and neurites of the CA1 region (Figure 8A). BAG3 was overexpressed at approximately four-fold of that of the control group and found largely limited to the hippocampal region (Figure 8B&C). BAG3 was found unaltered in the cerebellum, indicating the regional specificity of BAG3 overexpression (Supplemental Figure S3A&B).

**Figure 8.**
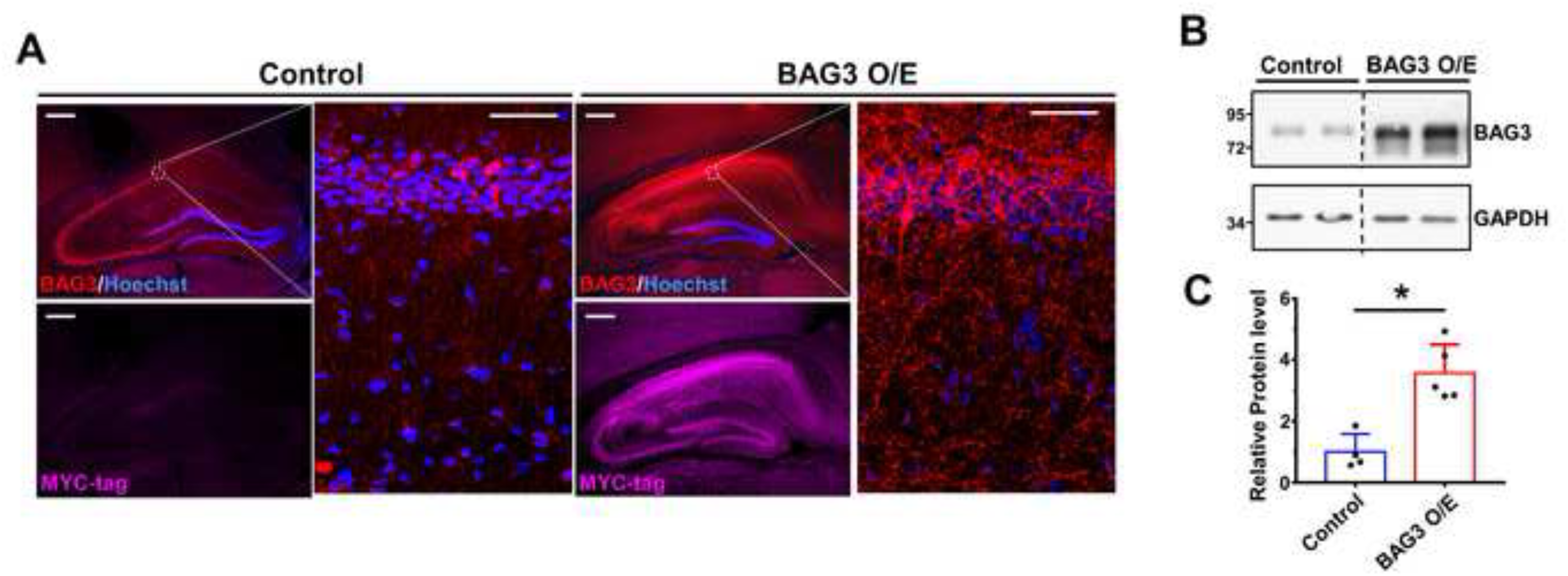
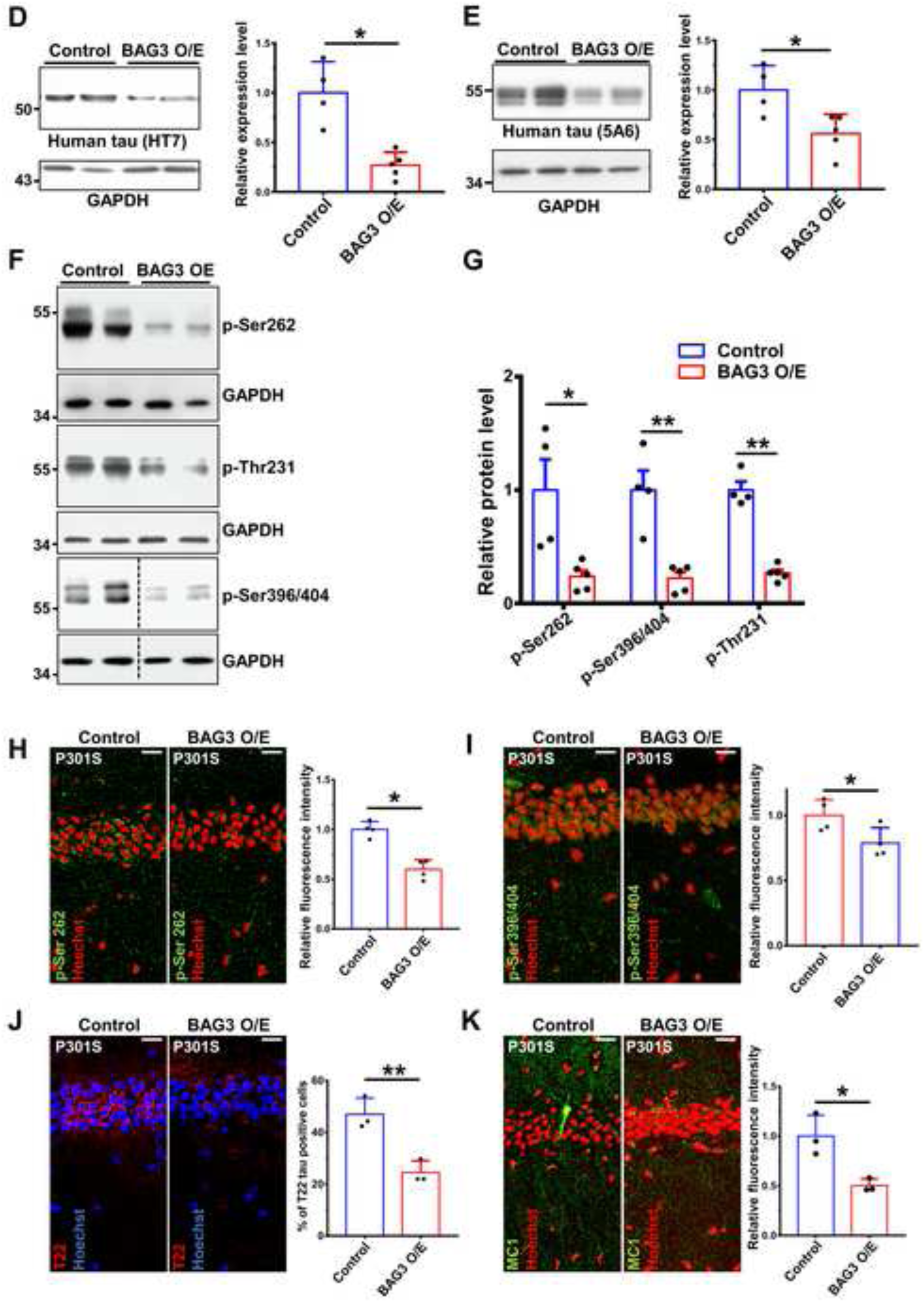
Overexpression of BAG3 in P301S mice in the hippocampus reduced the tau levels. Two-month-old P301S mice were intrahippocampally injected with AAV2/9-eGFP as control or AAV2/9-eGFP-hBAG3 (with a FLAG-Myc N terminal tag) for BAG3 overexpression (BAG3 O/E), and animals were collected at 6 months of age. (A) Images of the AAV injected brains were co-immunostained for BAG3 (red) and Myc tag (purple). The immunofluorescence images showed increased levels of BAG3 in the hippocampal region in the BAG3 O/E brain. Scale bars, 200μm. (B) Hippocampal lysates from control and BAG3 O/E brains were immunoblotted for BAG3. GAPDH is used as a loading control. (C) The graph shows the level of BAG3 normalized to GAPDH and relative to control. N=4 for control, n=5 for BAG3 O/E, Data are shown as mean ± SEM with Student’s t-test, *P<0.05. (D, E) Hippocampal lysates from control and BAG3 O/E brains were blot for total human tau (HT7 and 5A6). The graphs on the right show the quantitative analysis of tau normalized to GAPDH and relative to control. N=4 for control, n=5 for BAG3 O/E. (F) Representative blots of phosphorylated tau (p-Thr231, p-Ser262, and p-Ser396/Ser404) in the hippocampus lysate of control and BAG3 O/E brains. (G) Quantification of the levels of phosphorylated tau in control and BAG3 O/E hippocampal lysates. Data were normalized to the loading control GAPDH and then compared to controls, N=4 for control, n=5 for BAG3 O/E. (H, I) Phosphorylated tau immunofluorescence (p-Ser262 and p-Ser396/Ser404, Green) staining in the CA1 with corresponding intensity quantification relative to controls, N=4. (J) Oligomeric tau (T22 tau, purple) immunofluorescence staining in the CA1 with corresponding quantification, N=3. (K) Pre-tangle and tangle conformation tau (MC1 tau, green) immunofluorescence staining in the CA1 with corresponding quantification relative to controls, N=3. For all images, scale bars, 20μm and data are shown as mean ± SEM with unpaired Student’s t-test;. *, P<0.05; **, P<0.01.

Next, we investigated whether BAG3 over-expression modulated tau levels of P301S tau mice. Total human tau (recognized by the human tau specific antibodies HT7 (50) and 5A6 (51)) were significantly decreased in the BAG3 overexpression group (Figure 8D&E). The phosphorylated tau species including p-Ser262 tau, p-Thr231 tau and p-Ser396/404 tau were similarly significantly decreased in the BAG3 overexpression groups compared with the AAV control groups (Figure 8F&G), validating our *in vitro* studies. As expected, total human tau levels were not significantly changed in the cerebellum, showing that BAG3-mediated tau clearance is dependent on the regional overexpression of BAG3 (Supplemental Figure S3C&D). Quantitative IHC also showed p-Ser262 tau and p-Ser396/404 tau were significantly decreased in the CA1 region in the BAG3 overexpression brains (Fig 8H and I). Moreover, the percentage of oligomeric tau (recognized by T22 antibody (52)) positive cells was significantly reduced in the BAG3 overexpression group compared with the control group (Figure 8J). Staining with the conformation-dependent MC1 tau antibody (53) was also decreased in the CA1 in the BAG3 overexpression group (Figure 8K). These findings suggest that overexpression of BAG3 reduced the level of pathogenic tau species, indicating an essential role for BAG3 in tau clearance *in vivo*.

We further tested our hypothesis that BAG3 promotes the recruitment of Hrs to Rab35 *in vivo*. Co-immunostaining of Hrs and Rab35 in the P301S mouse brains showed significantly more Rab35 co-localized with Hrs in the CA1 region in the BAG3 overexpression group compared with the control group (Fig 9A-C). Furthermore, co-immunostaining of p-Ser396/404 tau with ESCRT III component, CHMP2B, showed co-localization of p-Ser396/404 tau with CHMP2B was significantly increased in the BAG3 overexpression group (Fig 9D-F). Finally, we examined the role of BAG3 in the synapse loss of P301S mice (47). IHC staining for MAP2 and PSD95 was used to visualize dendrites and postsynaptic compartments, respectively (54, 55). To quantify the density of MAP2 and PSD95, the images were reconstructed using the Imaris Surface Rendering Model. Our data showed the MAP2 and PSD95 density was significantly increased in the BAG3 overexpression group (Figure 10A-C). Overall, these findings suggest hippocampal overexpression of BAG3 alleviates tau pathology development in P301S tau transgenic mice.

**Figure 9.**
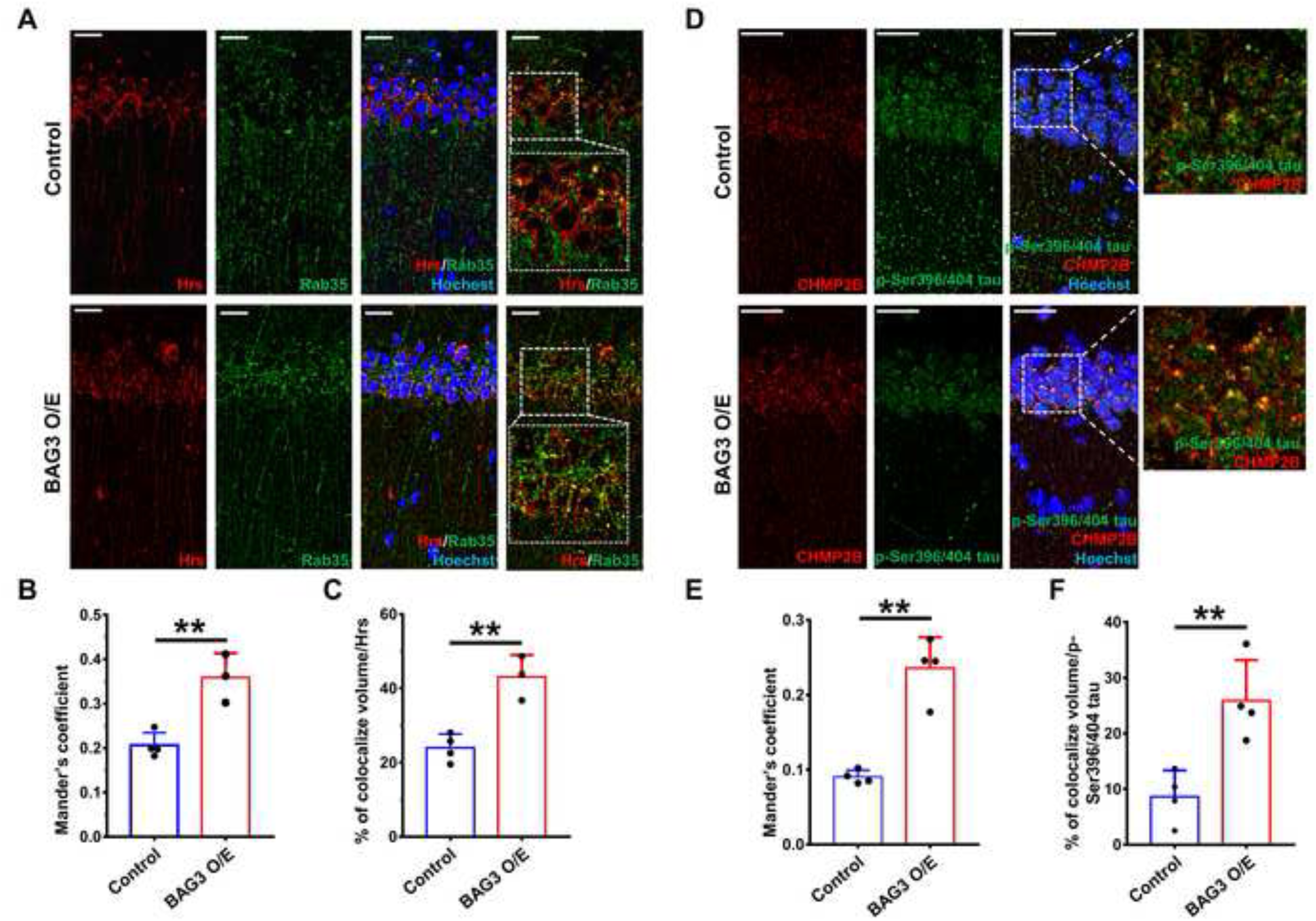
Overexpression of BAG3 in tau transgenic mice promotes Rab35-mediated recruitment of Hrs and tau sorting into the endocytic pathway. Two-month-old P301S tau mice were intrahippocampally injected with AAV2/9-GFP as control or AAV2/9-GFP-hBAG3 (with a FLAG-Myc N terminal tag) as BAG3 overexpression (BAG3 O/E) and collected at 6-month old. (A) Representative immunofluorescence staining and co-localization of Rab35 (Green) and Hrs (Red) in the CA1. (B) Quantification of co-occurring Rab35 in Hrs using Mander’s co-localization coefficient. (C) Quantification of the co-localization between Rab35 and Hrs based on volume. N=3 with unpaired Student’s t-test.. (D) Representative immunofluorescence staining of p-Ser396/404 tau (green) co-localization with CHMP2B (red) in the CA1. (E) Quantification of co-occurring of CHMP2B in p-Ser396/404 tau using Mander’s coefficient (F) Quantification of the co-localization between p-Ser396/404 tau and CHMP2B. All samples were counterstained with Hoechst 33342 (blue) to visualize the nuclei. Scale bars denote 20 μm. N=4 with unpaired Student’s t-test. *P<0.05, **P<0.01.

**Figure 10.**
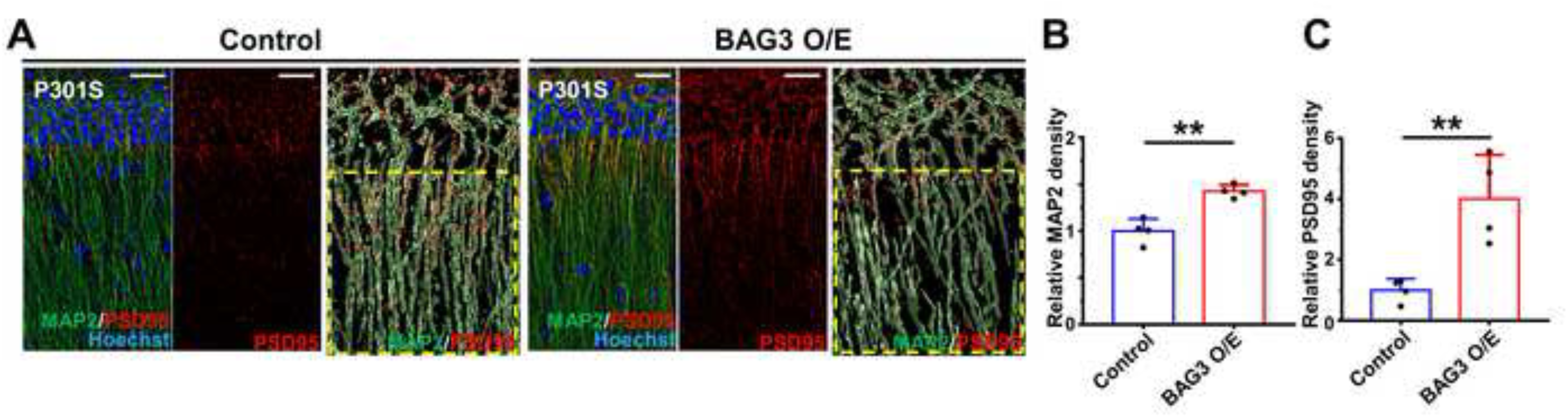
Overexpression of BAG3 in tau transgenic mice in the hippocampus increased the density of synapses and dendrites. (A) Representative immunofluorescence images of MAP2+ dendrites (green) and PSD95+ postsynaptic compartments (red). CA1 region of the hippocampus was imaged with Hoechst 33342 (blue) counterstaining for nuclei visualization. The border of each neuron was outlined using the ‘surface’ IMARIS function. Scale bars denote 20μm. (B&C) Relative density of MAP2 (B) and PSD95 (C) in the depicted area. The density of MAP2 or PSD95 were defined as the volume of fluorescence positive area/total area. Data are shown as mean ± SEM, N=4 using unpaired Student’s t-test, **, P<0.01.

## Discussion

Proteostatic dysregulation occurs during normal aging and could contribute to the pathogenesis of neurodegenerative diseases involving the pathological accumulation of protein, such as in AD which is defined by the presence of aggregates of abnormally phosphorylated tau (56–59). The cellular machinery involved in clearing toxic tau species is not fully understood, though evidence suggests that these pathways have strong therapeutic potential for the treatment and possible prevention of AD (60). Our lab has previously shown that BAG3 facilitates tau (21, 28). BAG3 is ubiquitously expressed and has numerous interacting partners ranging from synaptic proteins to many chaperone/co-chaperone proteins (28, 61, 62). The function of BAG3 is likely governed by the expression of cell-specific interactors and could be sensitive to over-expression, which may explain disparities in the BAG3 interactome literature (17, 63).

In this study, we assessed the endogenous neuronal BAG3 interactome and uncovered a novel mechanism of BAG3-mediated tau clearance through the endosome/lysosome pathway. We find that BAG3 interacts with TBC1D10B, the main regulator of Rab35, to control the trafficking and clearance of toxic tau species. Mutations in the different domains of BAG3 had little effect on tau clearance and TBC1D10B interaction apart from the L462P mutation, which impacts BAG3-HSP70 complexing (64), further supporting the importance of BAG3-HSP70 association in neuronal proteostasis. This BAG3-HSP70-TBC1D10B complex has important implications for the endosome/lysosome pathway. Since TBC1D10B inactivates Rab35 (24, 31), this complex implies BAG3-HSP70 monitoring of neuronal activities controlled by Rab35, such as presynaptic protein homeostasis and neuritic outgrowth, among others (31, 65, 66).

These findings support previous studies showing Rab35-mediated tau clearance (27) by demonstrating that BAG3 disinhibits Rab35 activity through its interaction with TBC1D10B and initiates downstream ESCRT activity through Hrs recruitment, MVB formation, and the trafficking of cargo such as tau species for degradation in an HSP70-dependent manner (27, 67, 68). Moreover, we provide the first evidence that upregulating BAG3 in disease-relevant regions of the brain *in vivo* ameliorates tau pathology by increasing engulfment by ESCRT machinery and rescues neuritic and synaptic morphology. We also show that the association of BAG3 with TBC1D10B is significantly less in AD cases compared to age-matched controls, which could contribute to a dysregulation of Rab35 and thus impaired tau clearance. Overall, the present study reports a novel BAG3-HSP70-TBC1D10B signaling axis for further study as a potent modulator of ESCRT-mediated endosomal protein clearance in aging and neurodegenerative diseases (Figure 11).

**Figure 11.**
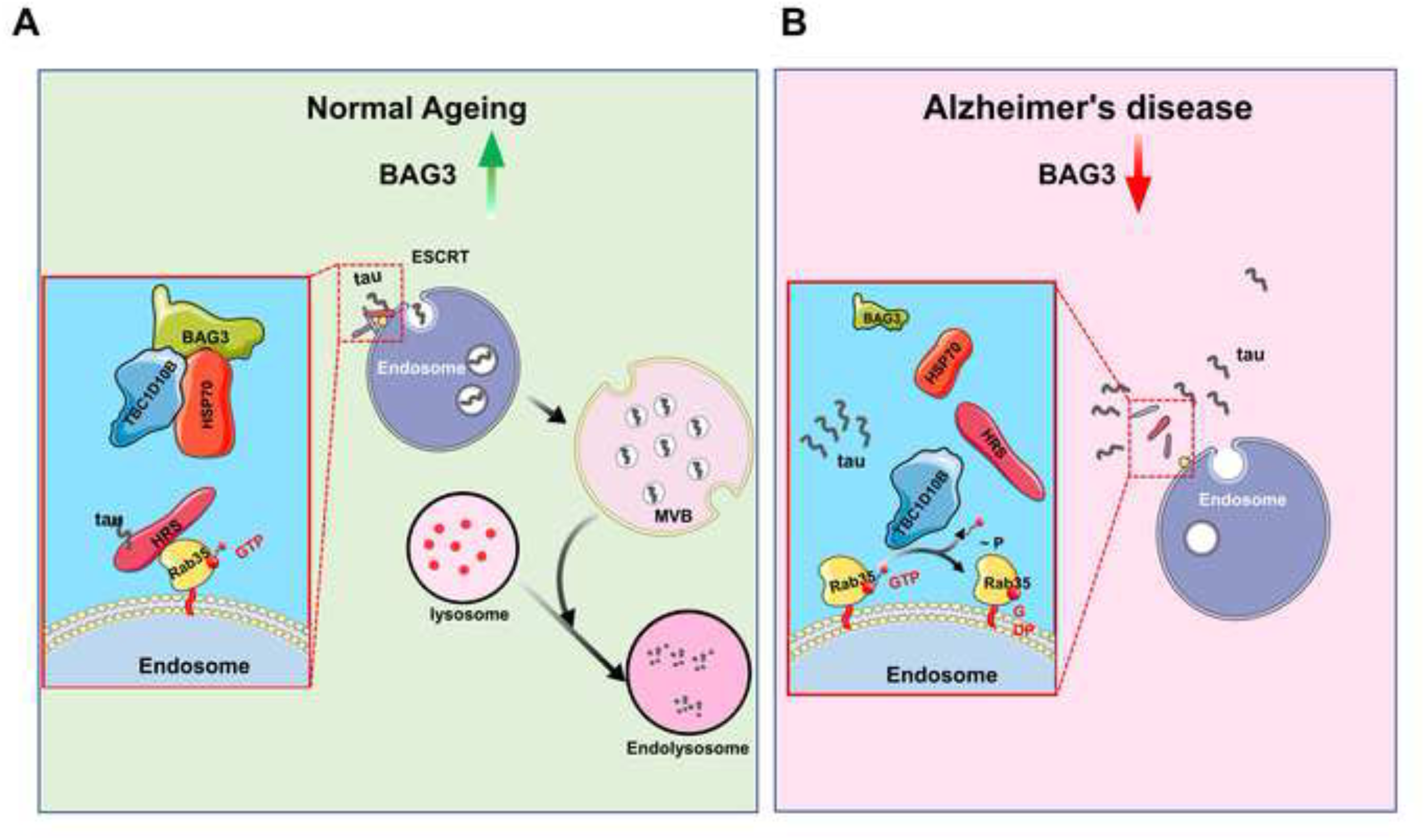
BAG3-TBC1D10B-Rab35 signaling axis regulates ESCRT and endosomal tau clearance. (A) During normal aging, BAG3 level increases, promoting the association of BAG3-HSP70-TBC1D10B. This prevents TBC1D10B from inactivating Rab35, leading to Hrs recruitment and initiation of ESCRT-mediated endosomal tau clearance. (B) In AD brains, however, this increase of BAG3 may be attenuated, which would release TBC1D10B from the BAG3-HSP70-TBC1D10B complex. Free TBC1D10B would thus be able to inactivate Rab35 and prevents endosomal clearance of tau, leading to accumulation over time.

## Acknowledgments

We thank Dr. C.Pröschel, University of Rochester for providing us with the pHUUG vector; Dr. J. Höfeld, University of Bonn for providing us with BAG3, L462P BAG3 and WAWA BAG3 in pcDNA; Dr. L.I. Binder for the gift of the MAP2 (AP14) and Tau5 antibodies [31, 32]; Dr. P. Davies for the gift of MC1 and phospho-Tau (Ser396/404) (PHF1) antibodies; Dr. P. Dolan for the gift of phospho-Tau (Ser262) antibody. Brain samples were obtained the UF Neuromedicine Human brain and Tissue bank (UF HBTB) at the University of Florida with informed consent of the patients or their relatives and the approval of the local institutional review boards. The UF HBTB is supported by the Florida ADRC (P30AG066506). This work was supported by National Institute of Health (NIH) grants R56 AG067739 and R01 NS098769. This research has been facilitated by a fee for service provided by the University of Rochester Mass Spectrometry Resource Laboratory and NIH instrument grant (S10OD021486).

## Author contributions

GJ contributed to the study conception and design, edited the manuscript, and provided funding. HL contributed to the study conception and design, performed experiments, analyzed the data, interpreted the experiments, wrote the manuscript. MT, CJ contributed to the study conception and design, performed experiments, analyzed the data and edited the manuscript. PG, GC, YC performed experiments, analyzed the data and edited the manuscript. SK assisted with data analysis and editing of the manuscript.

## Compliance with ethical standards

Conflict of interest: The authors declare that they have no conflicts of interest with the contents of this article.

Ethical approval All the work involving animals was reviewed and approved by the University Committee on Animal Research of the University of Rochester. (Protocol #:2007-023E&ER).

## Supplement 1

### Supplemental Methods and Materials

#### Reagents

##### Constructs

lentiviral vectors: shBAG3 (5’-AAGGTTCAGACCATCTTGGAA-3’) and BAG3-scrRNA (5’-CAGTCGCGTTTGCGACTGG-3’) in FG12 (with an H1 promoter) (1), or pHUUG with a U6 promoter (a generous gift of from Dr. C.Pröschel, University of Rochester) with and without GFP were used. shTBC1D10B (5’-GCTGTCTTAATTTGCCTTTGG-3’) and shRab35 (5’-CGATTGTGGTGATGTAGCTG-3’ targeting the ORF region and 5’-ATTTGTTAAGAGAATGCTCC-3’ targeting the 3’UTR region) were prepared in pHUUG. pcDNA3 TBC1D10B (NM_015527.3 (ORF sequence) in pcDNA3.1+/C-(K)-DYK (FLAG tagged)) was prepared by and purchased from Genscript (Genscript, NJ). Myc-Rab35 (plasmid#47433), Myc-Rab35 Q67L (plasmid #47434), eGFP-Rab35 (plasmid #49552), and pCS2 Hrs-RFP (plasmid #29685) were from Addgene. BAG3, WAWA BAG3 (2) (with W to A substitutions at positions 26 and 49), and L462P BAG3 in pcDNA (generous gifts from Dr. J. Höfeld, University of Bonn) were used as templates to clone into the lentiviral pCDH backbone (System Biosciences CD516-B2, CA) using Gibson assembly (New England Biolabs, #2611, MA). pCDH-GPG BAG3 was generated by mutation of the two IPV domains of BAG3 in pcDNA to GPG (with IPV to GPG substitution at position 95∼97 and 207∼209) and cloning into the pCDH backbone. pCDH-T4 tau-T2A-RFP was generated by clone T4 tau from pcDNA3.1 T4 tau (3) to replace the Puromycin resistant gene of the lentiviral pCDH backbone. pGEX-RBD35 was generated by cloning the Rab35-binding region of RUSC2 (RUN and SH3 domain-containing 2; amino acid residues 982– 1,199) from cDNA derived from mouse brain into pGEX-6P-2 (#GE28-9546-50, Millipore Sigma) at the BamH1 and Not1 sites. Lentiviral packaging vectors psPAX2 (#12260) and VSV-G (#12259) were from Addgene. AAV control virus (AAV9-SYN1-eGFP) and AAV BAG3 overexpression virus (AAV9-SYN1-GFP-T2A-BAG3 [with an N terminal FLAG-Myc tag]) were purchased from Vector Biolabs (Vector Biolabs, PA).

##### Antibodies

rabbit antibodies include: BAG3 (Proteintech, 10599-1-AP), TBC1D10B (Invitrogen, PA5-61832), V5 Tag (Cell Signaling Technology, #13202), Rab35 (Proteintech, 11329-2-AP), PSD95 (Cell Signaling Technology, #3450S), Hrs (Cell Signaling Technology, #15087), tau (Dako, A0024), phospho-Tau (Thr231)(AT180 Thermo Fisher OPA-03156), CHMP2B (Proteintech, 12527-I-AP), Normal Rabbit IgG (EMD Millipore 12-370). Mouse antibodies include: Hrs (Santa Cruz, sc-271455), Myc Tag (Cell Signaling Technology, #2276), FLAG Tag (Cell Signaling Technology, 8146), MAP2 (AP14, a gift from Dr. L.I. Binder (4, 5)), clathrin (Santa Cruz, sc-271178), MAP6 (Biolegend, 824701), MC1 (a gift from Dr. P. Davies), 5A6 (DSHB, Univ. Iowa), HT7 (ThermoFisher, MN1000B), tau5 (a gift from Dr L.I. Binder (4, 5)), T22 (Millipore Sigma, ABN454), phospho-Tau (Ser262) (12E8, a gift from Dr. P. Seubert), phospho-Tau (Ser396/404) (PHF1, a gift from Dr. P. Davies), GAPDH (Invitrogen, AM4300), Normal mouse IgG (EMD Millipore 12-371). Secondary antibodies include Alexa Fluor 594 donkey-anti-rabbit, Alexa Fluor 594 donkey-anti-mouse, Alexa Fluor 488 donkey-anti-mouse, Alexa Fluor 488 donkey-anti-rabbit, and Alexa Fluor 647 donkey-anti-mouse (Thermo Fisher Scientific). Conformation specific mouse anti-rabbit IgG (Cell Signaling Technology, #5127S). Anti-rabbit IgG HRP-conjugated Antibody (Bio-rad 5196-2504), Anti-mouse IgG HRP-conjugated Antibody (Bio-rad 5178-2504).

#### Animals

All mice and rats were maintained on a 12 h light/dark cycle with food and water available ad libitum. All procedures were approved by University Committee on Animal Research of the University of Rochester. Male P301S mice (PS19 line) were purchased from Jackson laboratories (Stock # 008169). There were 4-5 animals in each group for all quantitative analyses. All procedures were approved and performed in compliance with the University of Rochester guidelines for the care and use of laboratory animals.

To prepare samples for analyses, mice were deeply anesthetized with isoflurane and perfused with phosphate-buffered saline (PBS), followed by decapitation and rapid brain removal. The cortex and hippocampus of a half hemisphere were dissected out, collected, and quickly frozen for immunoblot analyses. The other hemisphere was fixed with 4% paraformaldehyde for 2 hours at 4°C, then cryoprotected by a transfer to 15% sucrose, followed by 30% sucrose solution in PBS until it sank in the latter. Each hemisphere was briefly dabbed with a lint-free wiper to remove excess sucrose and then rapidly frozen in cooled 2-methylbutane for 30 s on dry ice. Samples were kept at −80°C prior to sectioning and immunohistochemical analysis. All samples were stored at −80°C until further use.

#### Human brain samples

Human brain frontal cortex tissue were harvested from five individuals diagnosed with AD and five age-matched controls were obtained from the University of Florida Brain Bank. The average age of the individuals was 83.4 years for the AD patients and 81.6 years for aged controls. The aged controls were reported cognitive normal. Sex, post-mortem interval (PMI), Consortium to Establish a Registry for Alzheimer’s Disease (CERAD) and Thal score, Braak stage, for each case is documented. Brain tissues were paraffin embedded and sectioned at 5 µm.

| Sample | ID | Sex | Age (Years) | Thal | PMI (hours) | Braak | CERAD |
| --- | --- | --- | --- | --- | --- | --- | --- |
| Control 01 | 110001 | m | 90 | 1 | 14 | II | None |
| Control 02 | 110002 | m | 88 | 1 | 4 | II | None |
| Control 03 | 110003 | m | 72 | 0 | 4 | II | None |
| Control 04 | 110408 | m | 71 | 1 | 6 | I | None |
| Control 05 | A20-036 | f | 96 | 3 | 16 | II | Sparse |
| AD 01 | 110094 | m | 74 | 5 | 2 | V | Frequent |
| AD 02 | 110398 | f | 86 | 4 | 16 | VI | Frequent |
| AD 03 | 110373 | m | 83 | 5 | 18 | VI | Frequent |
| AD 04 | A19-008 | m | 81 | 5 | 4 | VI | Frequent |
| AD 05 | A20-004 | m | 84 | 4 | 16 | V | Frequent |

#### Stereotaxic surgeries

For BAG3 overexpression, 2-month-old P301S male mice received intrahippocampal injections of either AAV9-SYN1-eGFP or AAV9-SYN1-eGFP-2A-hBAG3 in both hemispheres and sacrificed 4 months post injection. For BAG3 and TBC1D10B colocalization analysis in Figure 2, an 8-month-old male wild type C57Bl/6 mouse (which had been intrahippocampally injected with AAV9 control virus at 6-month-old for another study). Animals received intrahippocampal AAV injections while under isoflurane anesthesia (1.75 % isoflurane in 30/70 % oxygen/nitrogen gas) using a Kopf stereotactic apparatus (6). Mice were secured using ear bars and a head holder. Ophthalmic ointment was applied to prevent drying of the eyes. Betadine was used to disinfect the scalp prior to incision with a scalpel. A 0.5-mm burr hole was drilled 1.8 mm caudal and 1.8 mm lateral from the bregma. A 33-GA needle was lowered 2 mm over 2 minutes. A Micro-1 microsyringe pump controller (World Precision Instruments) was used to inject 5 μL of AAVs using the convection-enhanced delivery (CED) method resulting in delivery of approximately 3×10^11^ infection particles/mL into each hippocampus as previously described (7). The needle was left in place for two additional minutes before it was slowly removed. The burr hole was filled with bone wax (Ethicon, Somerville, NJ), and the incision closed with 50 Dermalon sutures (Covidien, Mansfield, MA). Betadine and topical lidocaine were applied to the top of the suture to prevent infection and for analgesia, respectively.

#### Cell culture

Primary cortical neurons were prepared from rat embryos at E18 and cultured as previously described with some modifications (8). In brief, cerebral cortices were isolated from the embryonic rat brains, meninges were removed, then the cortices were transferred into trypsin-EDTA (0.05%) (Corning, MT25053CI). Digestion in trypsin occurred for 15-20 minutes in a 37°C water bath. Following gently trituration, neurons were plated at a density of 100,000 cells/cm^2^ for biochemical studies, and at a density of 10,000 cells/cm^2^ on coverslips for imaging. Both culture dishes and coverslips were coated with poly-D-lysine (Sigma, P6407). Neurons were grown up to DIV 22 in Neurobasal medium (Thermo Fisher Scientific) supplemented with 2% B27 (Thermo Fisher Scientific) and 2 mM GlutaMax (Thermo Fisher Scientific). Half of the media was replaced every 3-4 days. For lentiviral transduction, DIV7 neurons were treated with virus for 16-24 hours followed by a half medium change.

HEK293T cells (System Biosciences, #LV900A-1) were grown in DMEM medium supplemented with 10% fetal bovine serum (FBS), 2 mM GlutaMax and penicillin/ streptomycin on 60 mm dishes until 80% confluent. HEK cells were transfected with 2-3 μg of the designated construct or empty vector using the PolyJet (SignaGen Laboratories, SL100688). Cells were harvested 48hours post transfection for further analysis.

HEK293T cells were treated with 10 μM YM01 (9) (Sigma #SML0943) or DMSO as vehicle control at for 24 hours before collection for western blot analysis.

#### Creation of BAG3 knockout (KO) cell line by Crispr/Cas9 system

HEK293T cells were transfected with BAG3 gRNA plasmid (gRNA_eSpCas9-2A-GFP (PX458), GenScript). 20bp gRNA sequence was TCTGTCATGCCGCCACGTAA. Successful delivery of gRNA plasmid was confirmed by visualizing GFP expression. Cells were collected and resuspended in 1ml PBS with 2% FBS. Using fluorescence activated cell sorting (FACS), cells expressing GFP fluorescence were sorted into 96-well plate individually and allowed to grow for 14 days. Cell colonies were expanded, and the knockout of BAG3 was determined by immunoblotting. Tau (0N4R) expressing BAG3 null HEK293TN cells were generated by transducing BAG3 KO HEK293TN cells with T4 tau-T2A-RFP expressing lentivirus, followed by fluorescence activated cell sorting.

#### Lentivirus

To generate lentiviral particles, lentiviral vectors were co-transfected with viral packaging vectors into 70% confluent HEK293TN cells using PolyJet. Media were changed into DMEM containing 1% FetalClone-II serum (Hyclone, SH30066.03) 16h after transfection. The transfected HEK cells were then incubated at 33°C and 5% CO_2_ to slow their proliferation. Virus was collected at 64h after transfection and filtered through a 0.2 μm syringe filter to remove cell debris. Viral media was concentrated by ultracentrifugation at 4°C and resuspended in Neurobasal media. Concentrated virus was aliquoted, snap frozen, and stored at −80°C.

#### Mass Spectrometry Analysis

Rat primary cortical neurons transduced with either pHUUG-Scr or pHUUG-shBAG3 lentivirus at DIV16 were collected at DIV 22. BAG3 antibody (Proteintech, 10599-1-AP) was coupled to Dynabeads using the Dynabeads Antibody Coupling Kit (Thermo Fisher Scientific, Cat # 14311D). Cell lysates were immunoprecipitated with BAG3 antibody coupled beads. After washing, protein was eluted with a low pH elution buffer (0.1M citrate, pH 2.95). 1 M Tris (pH 8.8) was added immediately to the eluate to correct the pH and the samples were submitted to URMC Mass Spectrometry Resource Facility for mass spectrometry analysis.

Samples were run into a 4-12% SDS-PAGE gel to remove contaminants and create a ∼10mm length region. These regions were excised, cut into 1mm cubes, de-stained, then reduced and alkylated. Gel pieces were dehydrated and trypsinized at 37°C overnight. Peptides were extracted the next day, then dried down in a CentriVap concentrator (Labconco). Peptides were desalted, dried again, and reconstituted in 0.1% TFA. Peptides were injected onto a homemade 30 cm C18 column with 1.8 um beads (Sepax), with an Easy nLC-1000 HPLC (Thermo Fisher), connected to a Q Exactive Plus mass spectrometer (Thermo Fisher). The Q Exactive Plus was operated in data-dependent mode, with a full MS1 scan followed by 10 data-dependent MS2 scans.

Raw data was searched using the SEQUEST search engine within the Proteome Discoverer software platform, version 2.2 (Thermo Fisher Scientific), using the SwissProt Rattus norvegicus database. Label-free quantitation using Minora was used to determine relative protein abundance between samples. Percolator was used as the FDR calculator, filtering out peptides which had a q-value greater than 0.01. The identified proteins that showed a greater abundance in the Scr group compared to the shBAG3 group (PSM (Scr/shBAG3)>3) were selected and KEGG analysis was performed with String online platform (string-db.org).

#### Immunoblotting

Cell or tissue lysates were denatured in 1x SDS sample buffer at 100°C for 10 min before being loaded onto each lane of 10%-15% SDS-PAGE gels. After electrophoresis, proteins were transferred onto nitrocellulose membranes. Membranes were then blocked in TBS-T (0.1% Tween-20) containing 5% non-fat dry milk for 1h at room temperature. Primary antibodies were diluted in blocking solutions followed by incubation at 4°C overnight. The next day, membranes were further incubated with secondary antibody for 1h at room temperature. After thoroughly washing, membranes were visualized by enhanced chemiluminescence and images captured using the KwikQuant™ Imager (Kindle Biosciences, LLC). The intensity of each band was quantified using Image Studio Lite (Li-Cor). GAPDH was used as loading controls. Treatments were then normalized to their corresponding control sample and expressed as a fold difference above the control.

#### Immunohistochemical staining

Brain slices (30 µm) were prepared from mouse brains fixed with 4% paraformaldehyde using a sliding microtome (VWR 89428-70) with a freezing stage (PhysiTemp Instruments BFS-3MP/PTU-3). The brain sections were washed three times using 0.15 M phosphate buffer (0.05M NaH_2_PO_4_, 0.1M Na_2_HPO_4_) to remove the cryoprotectant. Sections were mounted on poly-D-lysine coated slides (Thermo Fisher), antigen retrieval was performed, and the slides were blocked with PBS containing 5% BSA and 0.1% tween 20 for 1 h at room temperature. The sections were incubated with primary antibody in 5% BSA in PBS overnight at 4°C. The next day, slices were incubated for 1 h at room temperature with Alexa Fluor™-conjugated secondary antibody including Alexa Fluor™ 594 donkey-anti-rabbit, Alexa Fluor™ 488 donkey-anti-rabbit, or Alexa Fluor™ 647 donkey-anti-rabbit (Thermo Fisher Scientific). Alternatively, they were labeled using the MOM kit (BMK-2202, Vector laboratories), followed by three washes with PBS and labeling with Streptavidin Alexa Fluor™ 488 or 647 (Thermo Fisher Scientific). The brain sections were coverslipped with ProLong Diamond Antifade Mountant (Thermo Fisher Scientific, P36961). The slides were imaged using a Nikon A1R HD laser scanning confocal microscope (Nikon). The brain sections were imaged at CA1 of the hippocampus with Nikon ECLIPSE Ti2 and recorded by NIS-Elements (Version 5.11) software. Resulting images were pseudocolored for illustration purposes.

#### Immunofluorescence

Neurons grown on coverslips were rinsed with PBS twice and followed by fixing in PBS containing 4% paraformaldehyde and 4% sucrose for 5 minutes at room temperature. Then, neurons were permeabilized in PBS containing 0.25% Triton X-100 and were blocked with PBS containing 5% BSA and 0.3 M glycine. Primary antibodies were diluted in blocking solution, added to the coverslips, and incubated on a shaker at 4°C overnight. Alexa Fluor 488/594/647 conjugated secondary antibodies were diluted in blocking solution and incubated with neurons for 1 hour at room temperature. Coverslips were counterstained with Hoechst 33342 and mounted with ProLong Diamond Antifade Mountant. Images were acquired on a Nikon A1R HD scanning confocal microscope using a 40x objective with a 2x optical magnification. Resulting images were pseudocolored for illustration purposes.

#### Live cell imaging

BAG3 null HEK293TN cells on 25mm coverslips were transfected with control vector or BAG3 expression vector together with GFP-Rab35 and Hrs-RFP. The coverslips were placed in an Attofluor™ Cell Chamber (#A7816, Thermo Fisher Scientific) with Krebs-Ringer Solution at 36 hours post transfection. During imaging, cells were placed inside a heating chamber (TIPA plate adapter for Nikon TIA) at 37 °C. Live cell imaging of fluorescent proteins was performed using a Nikon A1R HD laser scanning confocal microscope and images were acquired with a 60x Oil objective. Images obtained by Nikon ECLIPSE Ti2 and recorded by NIS-Elments (Nikon, Version 5.11) software at a rate of 6 frame per minute for 10 minutes.

Live cell image stacks were analyzed using Imaris Cell Imaging Software (Bitplane, Switzerland). Briefly, kymographs were generated using the function of swap time and z, and the rendered images were rotate 45 degree. The mobility of Hrs-RFP was determined by using the spots module in Imaris. First spots were generated for the Hrs-RFP range of 1∼2μm in diameter. Next, the spots were tracked over 10 minutes and the track of the spots were recorded. Finally, the total length of the different tracks was binned every 0.1 μm. Over 10 cells for each group were used for analysis.

#### Immunoprecipitation

Cells were lysed in ice-cold lysis buffer (50 mM Tris, 150 mM NaCl, 0.4% NP-40, 1 mM EDTA, 1 mM EGTA, pH7.4) supplemented with protease inhibitor cocktails and phosphatase inhibitors. Cell lysates were briefly sonicated then centrifuged at 13,200 rpm for 10 minutes at 4°C. Cleared supernatants (500 μg) were mixed with 2 μg normal rabbit/mouse IgG control or primary antibody, followed by incubation at 4°C for 24h. Antibody/antigen mixture was incubated with Dynabeads M-280 sheep anti-rabbit or mouse IgG (Thermo Fisher Scientific) for 6h at 4°C on the following day. An aliquot of protein lysate was saved for input control. After thoroughly washing the beads, bound fractions were eluted in sample loading buffer by boiling at 100°C for 10 minutes. Proteins were then resolved by SDS-PAGE. To reduce the background of heavy chain and light chain from IgG and antibodies, a conformation-specific mouse anti-rabbit IgG, a secondary rabbit antibody, was used following the precipitation of TBC1D10B.

#### Rab35 activity assay

The Rab35 activity assay was performed as previously described (10). Briefly, pGEX (GST only) or pGEX-RBD35 were transformed into BL21 E. coli, and GST and GST-RBD35 protein expression was induced with IPTG, followed by purification using glutathione S-transferase beads (Ge healthcare, #17-5132). The proteins were eluted and concentrated with Amicon Centrifugal Filter Units 10K (GST) and 50K (GST-RBD35) (Millipore) to remove excessive glutathione. Cell lysates from 10 cm dishes were collected and incubated with 1.25 μg of GST or GST-RBD35 recombinant protein, followed by precipitation with glutathione S-transferase beads. The precipitated samples were analyzed by immunoblotting and 1∼4% of the cell lysates were used as input controls.

#### Image analysis

Immunofluorescence images were opened and processed with combination of Image J and Imaris. Regions of interests were selected around either the soma, or a line (line width equals 120 pixels) was drawn along processes, followed by straightening with Imaris. Line scans were generated using the plot profile tool in Fiji (line width of one pixel). Intensity values for a given marker were normalized by subtracting the minimum background and dividing the dataset by its maximum.

Colocalization analysis was done using Imaris. Specifically, regions of interest were selected at the molecular layer in the CA1 region of the hippocampus (Figure 2B, Figure 8H, I, J and K, Figure 9 A, D, F and Figure 10A) or along processes with a line width of 200 pixels, followed by straightening (Fig 2C-E, Figure 6A, Figure 7A and D). Co-localization module was used, and different fluorescence channels were thresholded to generate a new colocalization channel. Mander’s colocalization coefficients were used to quantify the overlapping of fluorescence intensity(11).

Western blot images were turned into Black and white with Photoshop software (Adobe, Ver, CC). And the grey value was analyzed with image studio lite (LI-COR Biosciences, Ver 5.0).

#### Statistical analysis

All image measurements were obtained from the raw data. GraphPad Prism was used to plot graphs and perform statistical analysis. The statistical tests used are denoted in each Figure legend, and statistical significance was defined as *p < 0.05 and **p < 0.01. For the live-cell imaging analysis, the difference of Hrs mobility curves (Figure 6D) was transformed into an accumulation curve and then analyzed using the Kolmogorov–Smirnov test. All immunoprecipitation data are repeated at least twice.

**Supplemental Table 1. BAG3 associated proteins identified by immunoprecipitation-mass spectrometry.**

## Supplementary Figures

**Supplementary Figure S1.**
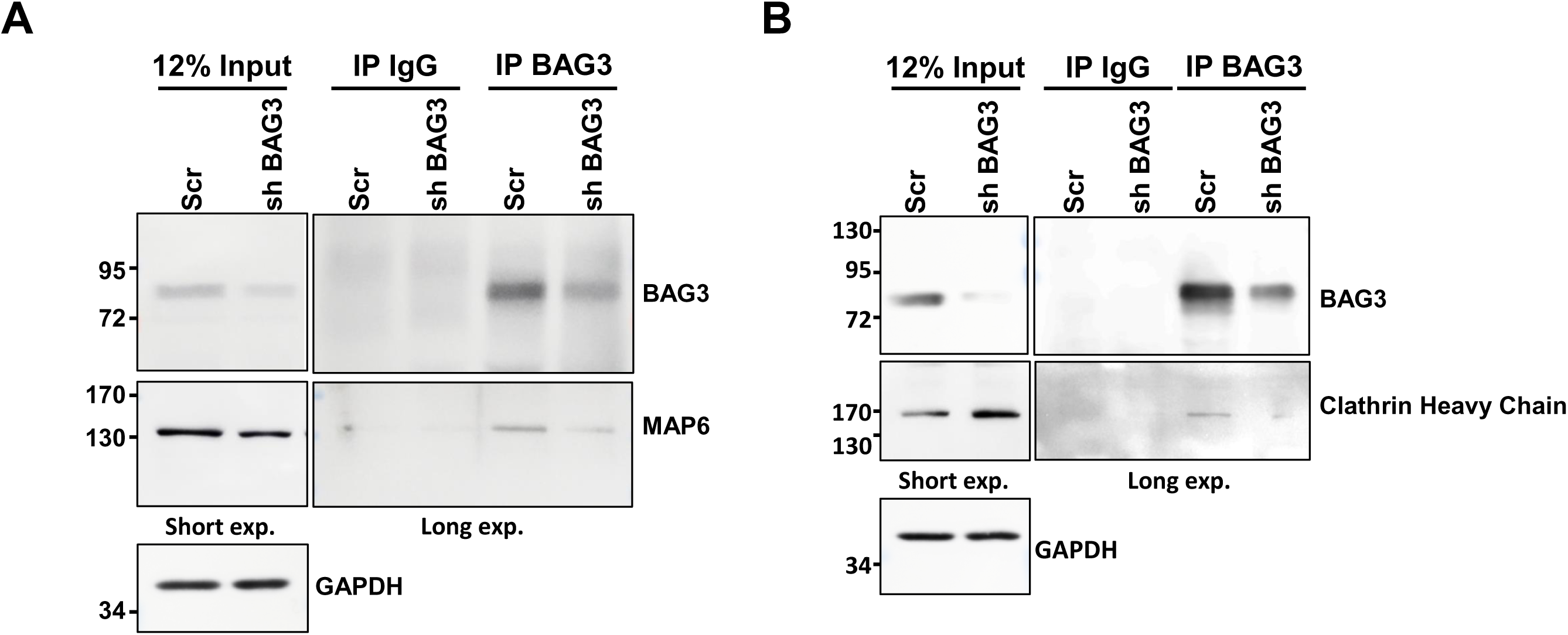
Verification of immunoprecipitation-mass spectrometry data. Cell lysates from rat neurons transduced with lentivirus expressing scrambled or shBAG3 shRNA at DIV16 were collected at DIV 22. MAP6 and clathrin were selected from the list of proteins that co-immunoprecipitated with BAG3 that were identified by mass spectrometry for verification. (a) Cell lysates were immunoprecipitated with an anti-BAG3 antibody and probed for the presence of MAP6. (b) Cell lysates were immunoprecipitated with an anti-BAG3 antibody and probed for the presence of clathrin. The same amount of rabbit IgG was used to verify the specificity of the immunoprecipitation. A fraction of cell lysate was used as input control. GAPDH is used as a loading control. the positions at which molecular weight markers (kDa) migrated are indicated at the left.

**Supplementary Figure S2.**
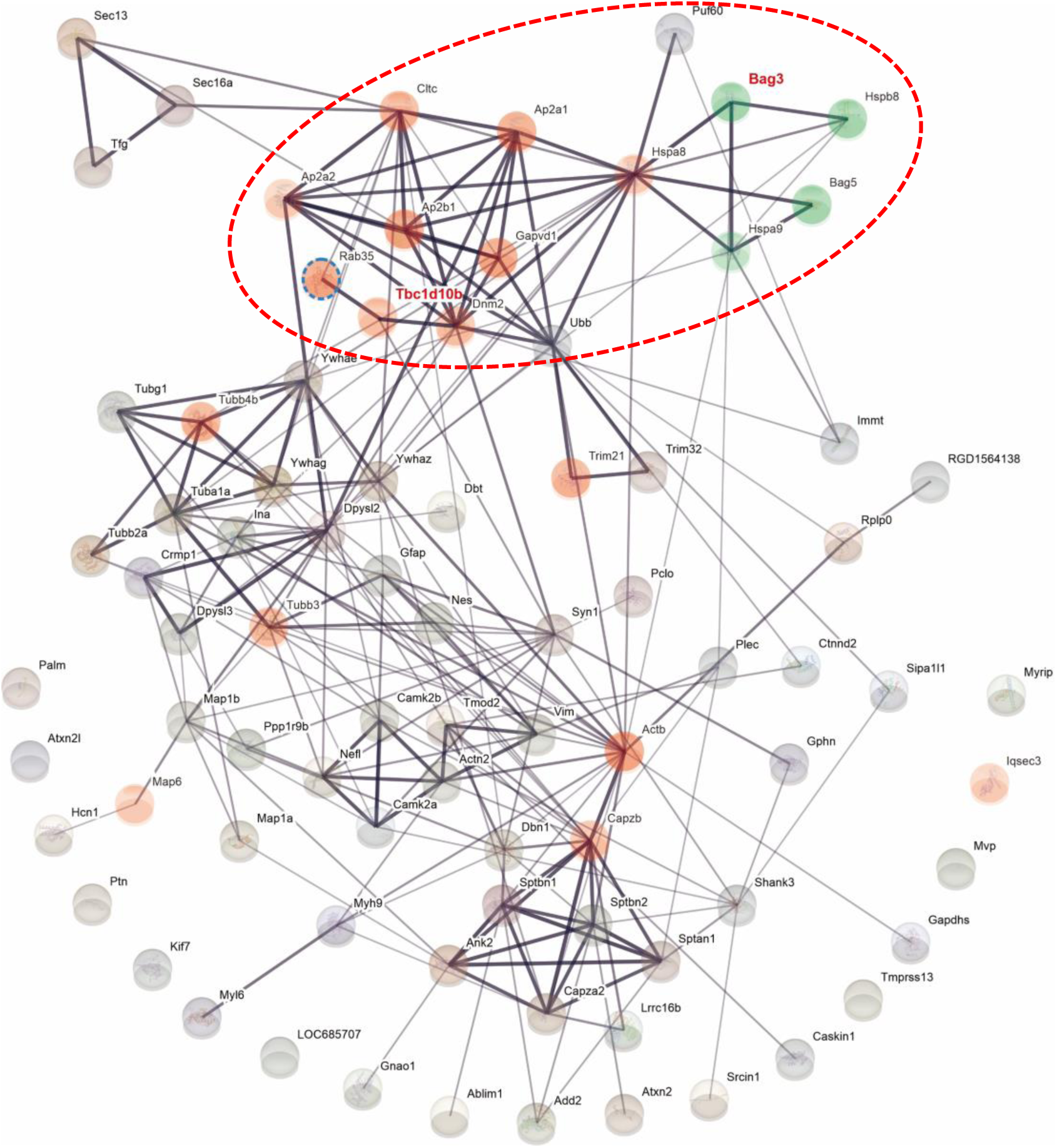
STRING analysis of predicted protein-protein interactions for the identified BAG3 associated proteins based on confidence. STRING analysis reveals that BAG3 associated with a diverse set of proteins which influence on endocytosis. STRING analysis reveals that BAG3 associated with a diverse set of proteins which influence on endocytosis pathway. Magnified image of Figure 1B.

**Supplementary Figure S3.**
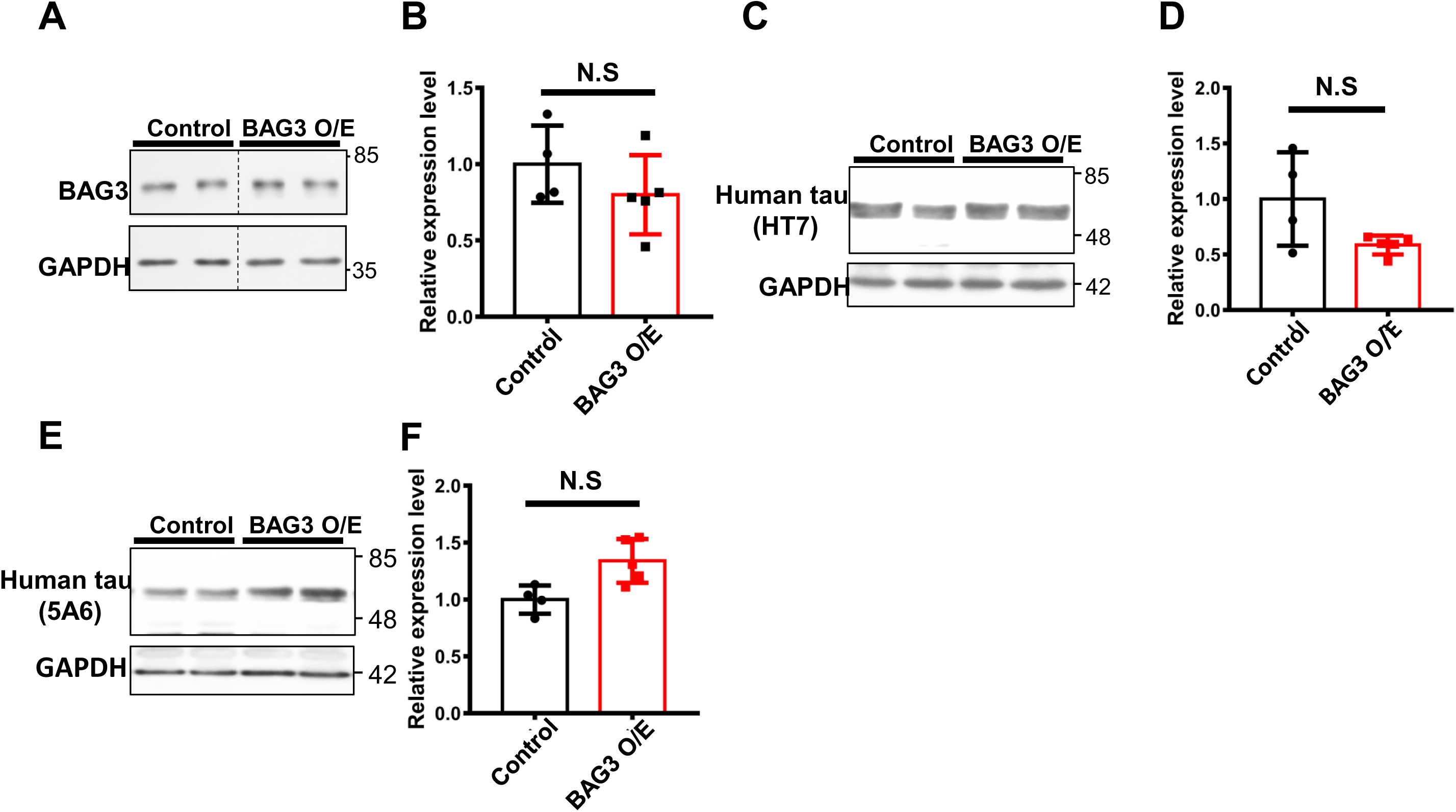
Overexpression of BAG3 in the hippocampus doesn’t affect the tau levels in the cerebellum. Two-month-old P301S mice were intrahippocampal injected with AAV2/9-GFP as control or AAV2/9-GFP-BAG3-MYC as BAG3 overexpression (BAG3 O/E) and were collected at 6-month old. (a) Representative immunoblots of BAG3 for cerebellum lysate of control and BAG3 O/E brains. (b) The graph shows the quantitative analysis of BAG3 to GAPDH and relative to control. (c, e) Representative blot of total tau (HT5 and 5A6) for cerebellum lysates of control and BAG3 O/E brains. (d, f) The graphs show the quantitative analysis of tau to GAPDH and relative to control. N=4 for control, n=5 for BAG3 O/E. Data are shown as mean ± SEM. Statistical analysis was performed using the student t-test. NS, not significant.

